# Beyond the Default: Optimizing Molecular Networking with arteMIS

**DOI:** 10.64898/2026.08.17.745252

**Authors:** Laura Rosina Torres-Ortega, Esteban Charria-Girón, Florian Huber, Matteo Simone, Margherita Sosio, Justin J.J. van der Hooft

## Abstract

Metabolomics uses tandem mass spectrometry (MS/MS) data to gain structural insights of small molecules that play biological roles, generating datasets whose size and complexity demand systematic organisation. Molecular networking addresses this by representing MS/MS spectra as nodes and their pairwise similarity as edges, but its output is critically sensitive to user-defined parameters: similarity score cut-off, maximum component size, maximum links and minimum matching peaks. These parameters are routinely left at default values, which can either collapse interpretable molecular families into entangled “hairballs” or fragment them into disconnected singletons. In the absence of ground truth, no standardised framework exists to evaluate molecular networks or to assess whether their connections are robust to run-to-run variability present in metabolomic experiments. Here, we introduce arteMIS (Accelerated Ranking and Tuning using Multi-metric Interpretability across Scores), a framework for systematic parameter optimisation that uses Latin Hypercube Sampling to efficiently cover the four-dimensional parameter space and ranks candidate networks through a user-tuneable composite Z-score, combining topology-and chemistry-based metrics. This framework supports three complementary modes: global, seed, and target-class, adapting optimisation to fully unannotated datasets, curated subset of reference features or class-focused discovery, respectively. Benchmarking across four spectral libraries (∼600 to ∼13,000 spectra) and four scoring methods (Cosine, Modified Cosine, Spec2Vec, MS2DeepScore), we provide practical guidance for parameter selection as a function of scoring method and dataset size and show that optimal settings do not transfer between them. Top-ranked arteMIS configurations matched or outperformed GNPS defaults in chemistry and topology metrics and produced networks with higher edge-stability under subsampling. Applied to actinobacteria and fungal samples, arteMIS rescued structurally meaningful families that remained fragmented under default settings. We conclude that arteMIS reframes molecular network construction from a default-driven step into a task-customisable optimisation.

Audience: metabolomics community, computational mass spectrometry, natural products

## Introduction

Mass spectrometry (MS) coupled with liquid chromatography (LC) is currently the most widely used platform for characterising the chemical diversity of small molecules within complex biological systems^1^. Tandem Mass Spectrometry (MS/MS) captures structural insights that can be used for molecular annotation under the hypothesis that similar fragmentation patterns imply chemical similarity^2,3^. As modern instruments now easily produce millions of MS/MS spectra across studies, computational approaches have become essential to organise these data and translate spectral patterns into structural hypotheses^3^.

Molecular networking (MN) has been introduced and further developed to address this challenge by organising, visualising, and propagating spectral annotations at scale^4,5^. In these molecular networks, nodes represent MS/MS spectra or MS/MS-derived metabolite features, and edges reflect the spectral similarity scores (e.g., Modified Cosine) above a user-defined cut-off threshold. Beyond organisation in a network graph, MN enables propagation of spectral annotations across subnetworks or so-called molecular families (MFs) exceeding direct library matching alone^6–8^. Implemented most broadly through the GNPS platform^4^, with recent advances like Feature Based Molecular Networking^9^ (FBMN) and Ion Identity Molecular Networking^10^ (IIMN). These approaches have become widely adopted for exploring complex natural product mixtures in microbiomes, plants, microbes, and marine organisms^11,12^, with applications including natural product dereplication and discovery^13–17^, pathway reconstruction^18^, gene cluster annotation^19,20^, drug metabolism^21–23^, among others^24^.

Despite their successes, current MN approaches face challenges that limit their reproducibility and interpretability. MN construction involves two key steps: (i) pairwise mass spectral similarity computation and (ii) network construction based on user-defined parameters. Although substantial efforts have focused on improving similarity metrics for the first step, the impact of network construction parameters has been largely overlooked^25^. Small variations in parameters, such as similarity threshold, maximum component size and maximum links, among others, can drastically reshape network topology, collapsing interpretable molecular families into overconnected “hairballs” or fragmenting them into disconnected singletons. A global similarity cut-off, for instance, is unlikely to perform equally well across all chemical families, some of which produce sparser or overlapping fragmentation patterns^26,27^.

Moreover, because similarity scores fundamentally determine network connectivity, parameter choices cannot be assumed to transfer across different scoring methods; beyond conventional cosine-based metrics, machine-learning-based approaches such as Spec2Vec^28^ and MS2DeepScore^29^ can capture molecule relationships under multiple chemical modifications; however, their behaviour under typical MN construction processes is poorly characterised, and practical guidance for parameter selection across datasets and scoring methods is largely absent.

Evaluating molecular networks itself is a fundamental challenge. Unlike supervised tasks, MN lacks ground-truth labels for edges: spectrally related molecules may share substructures, adducts, charge variants, in-source fragments, or be isomers, and many nodes remain unannotated by library matching. Therefore, network evaluation is done indirectly, balancing chemically plausible connections against global topology. Recent efforts, including Nth-neighbour-style metrics, network-level consistency and accuracy measures^25^, represent important steps toward quantitative assessment. However, these metrics also have important limitations. For example, the Nth-neighbour-style can assign high performance to a “hairball” network in which most spectra are connected. Moreover, the consistency measurement includes singletons in the calculation, which can inflate performance. These limitations highlight the complexity of evaluating molecular networking at the network level^30^.

Here, we introduce arteMIS, a systematic and customisable framework to optimise MN construction and evaluation. Rather than relying on default parameters or manual tuning, we sample the parameter space: similarity cut-off, minimum matching peaks, maximum component size, and maximum links, using Latin Hypercube Sampling (LHS), ensuring a broad and efficient coverage across a high-dimensional space. For each sampled configuration, we compute a set of complementary metrics evaluating both topology and chemistry on the constructed network.

To make this framework applicable in different experimental settings, arteMIS provides three configuration modes that differ in the origin of the chemical annotations used and in what the composite score is optimised for. In global-mode, annotations are available for every feature, either directly from spectral library metadata or by using tools like MS2Query^30^ that predict chemical classification terms for each spectrum, and networks are ranked by the overall chemical metrics: intra-molecular family similarity and NPClassifier^31^ consistency, representing a direct use with no prior manual curation. In seed-mode, only a user-subset of high confidence annotated features (e.g., features from spectral library hits or in-house standards) is used to score chemistry-based metrics, thereby avoiding the use and propagation of uncertain MS2Query predictions. In target-class mode, the aim is to cluster a specific compound class as accurately as possible in the molecular network (e.g., rescuing a fragmented family of specialised metabolites). Together, these modes apply the same optimisation strategy to fully unannotated datasets, curated reference subsets or class-focused discovery efforts within a single framework.

We evaluated parameter sensitivity across four library datasets of increasing size and assessed network robustness through a node-resampling strategy, repeatedly subsampling 85% of nodes to quantify edge stability and node isolation probability. Finally, we apply this workflow to case studies on bacterial and fungal specialized metabolites, demonstrating that optimised parameters rescue chemically coherent connections lost under default settings and molecular relationships that would otherwise remain hidden.

## Results and Discussion

### 1. Parameter influence of metrics across datasets and scores

Molecular network structure and chemical interpretability depend strongly on user-defined parameters, yet practical guidance for selecting values that preserve chemically meaningful groupings without over-fragmenting the network remains limited. To assess parameter effects, we selected four natural product (NP)-relevant datasets containing reference data of different and increasing spectral size (ranging from ∼600 to ∼13,000 spectra) and four different spectral similarity scores, and generated 100 networks per dataset per similarity score (1600 networks in total) using Latin Hypercube Sampling (LHS) across four commonly used parameters: similarity cut-off, matching peaks, maximum component size, and maximum links (for full description and ranges see Methods – Table 5 and 6). Each network was evaluated using three topology-based metrics (Coefficient of variation of node degree, Gini coefficient and % of isolated nodes; see Methods) and two chemistry-based metrics (intra-similarity and NPClassifier^31^-based consistency; Table 1). The running times and discrepancy are reported at Supplementary Table 1.

**Table 1.** Topology and chemistry-based descriptive metrics.

| Name | Concept | Type |
| --- | --- | --- |
| Coefficient of variation of node degree (CV_degree) | Calculates the coefficient of variation (std / mean) of node degrees. Higher values represent more disparity in the network, not modular. | Topology |
| Gini-coefficient | The Gini coefficient of connected molecular families. Lower values represent more modular families while higher values imply that one or few molecular families dominate the network. | Topology |
| % Isolated nodes | % of isolated nodes present in the network. | Topology |
| Intra-similarity | Mean pairwise Tanimoto similarity within components (Morgan fingerprints). Higher values represent molecular families containing more similar neighbours. | Chemistry |
| Consistency | Fraction of nodes whose members share the predominant NPClassifier annotation with a molecular family of purity $\geq 0.7$. Higher values represent molecular networks with more molecular families with the same class label. | Chemistry |

We note that we initially included adjusted mutual information (AMI) as a chemistry metric but excluded it after finding that it penalises the biologically expected splitting of a single chemical class across multiple pure molecular families; rationale and comparison are detailed in Methods – Evaluation metrics section, the actual metrics used, and their descriptions kept are listed in Table 1.

Across all four scores and datasets, raising the similarity cut-off value increases chemical metrics (especially consistency), but at the cost of more isolated nodes (Spearman correlations shown in Figure 1. and Supplementary Figures 1-3 for the other scores), a central known trade-off in network construction^32^. Maximum component size lowers intra-similarity generally, since a higher cap lets chemically distant nodes join the same family. Matching peaks (for the cosine-based scores – see for the machine learning-based scores further below) behaves like cut-off in the smaller datasets but not in the larger ones, where its effect on topology reverses (Gini coefficient increases and intra-similarity decreases). Maximum links affects only the degree distribution (CV of node degree), and only once datasets are large enough to reach the cap.

**Figure 1.**
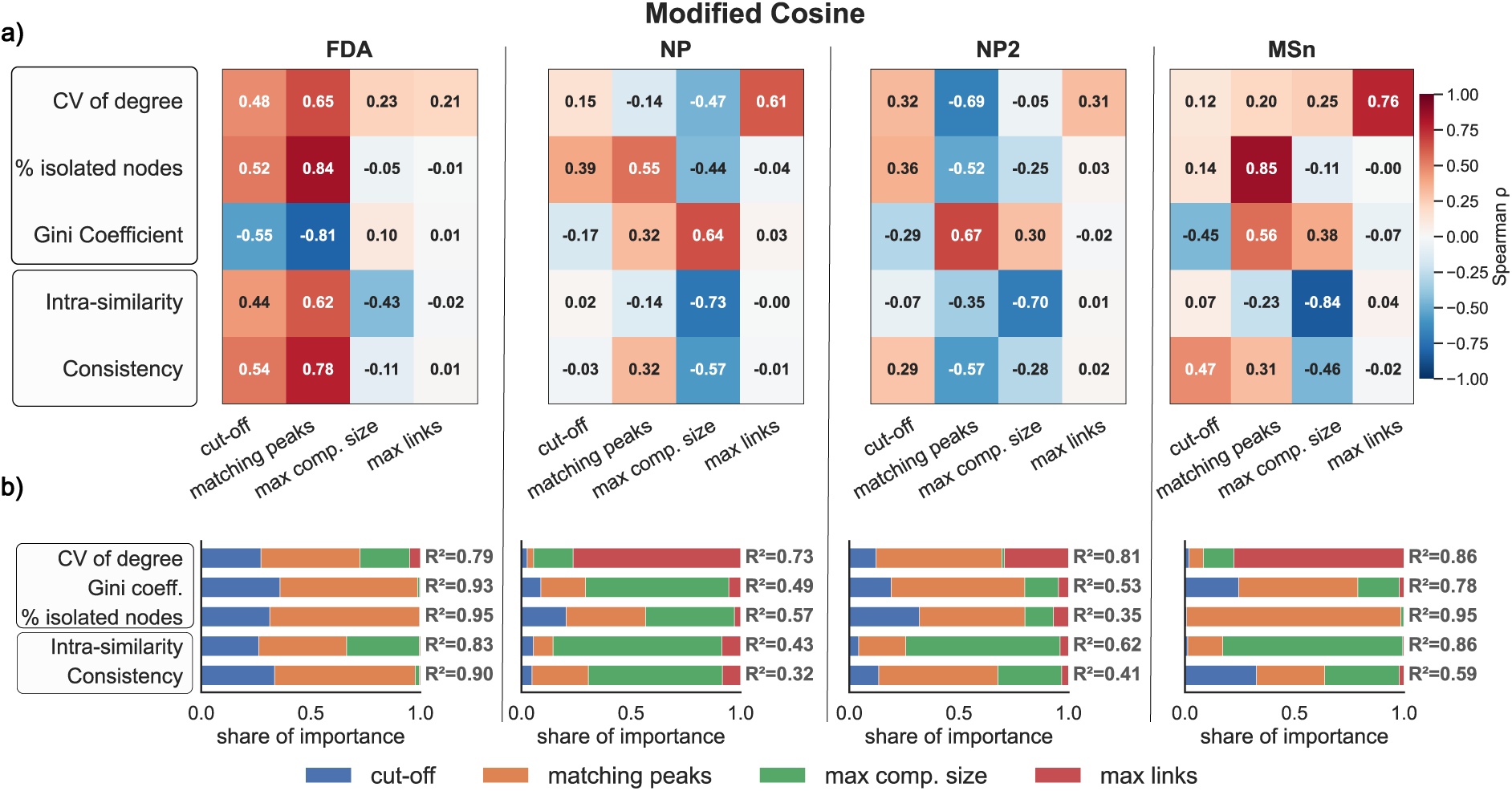
Parameter sensitivity of modified cosine across datasets where FDA-dataset contains 606 spectra, NP-dataset contains 1226, NP2-dataset contains 7250 spectra and MSn-COCONUT contains 12929 spectra. (a) Spearman correlations between network parameters and evaluation metrics for all four datasets. (b) Random-forest feature importances per dataset.

Although Spearman correlations reveal the direction (positive or negative) of each parameter’s influence on the metrics, it does not reveal its relative importance. To quantify how much each parameter contributes, and capture non-linear effects the correlation misses, we trained a random-forest per metric and examined feature importances. The results confirm the effects and add their magnitude: similarity cut-off and matching peaks account for most of the explained variance in the chemistry and isolation metrics, while maximum component size dominates intra-similarity (Figure 1b). The four parameters explain the metrics well in both the smallest and largest library datasets (R² up to ∼0.90) but capture less of the variation in the mid-sized natural-product sets, as shown in Figure 1b (R² = 0.32–0.80).

The parameters effects and their magnitudes do not transfer across scores. For cosine (Supplementary Figure 4), similarity cut-off and minimal number of matching peaks drives the importance across datasets. For Spec2Vec, cut-off alone drives nearly everything (see Supplementary Figure 5); maximum component size adds an effect only in the larger datasets; while for MS2DeepScore maximum component size drives most metrics across every dataset size and is the largest contributor to intra-similarity (Supplementary Figure 6). Since the parameters that matter change with the spectral similarity score, settings tuned on one similarity score cannot be reused on another.

We observe a chemistry–topology trade-off between metrics: chemically meaningful networks come at the price of fragmentation (Figure 2 (a)). Consistency rises with the fraction of isolated nodes across every score and dataset. MS2DeepScore is the exception worth noting; it reaches high consistency at lower isolation, particularly for the NP and MSn datasets, occupying a desirable trade-off region that the other similarity scores did not reach (Figure 2 (a) in pink).

**Figure 2.**
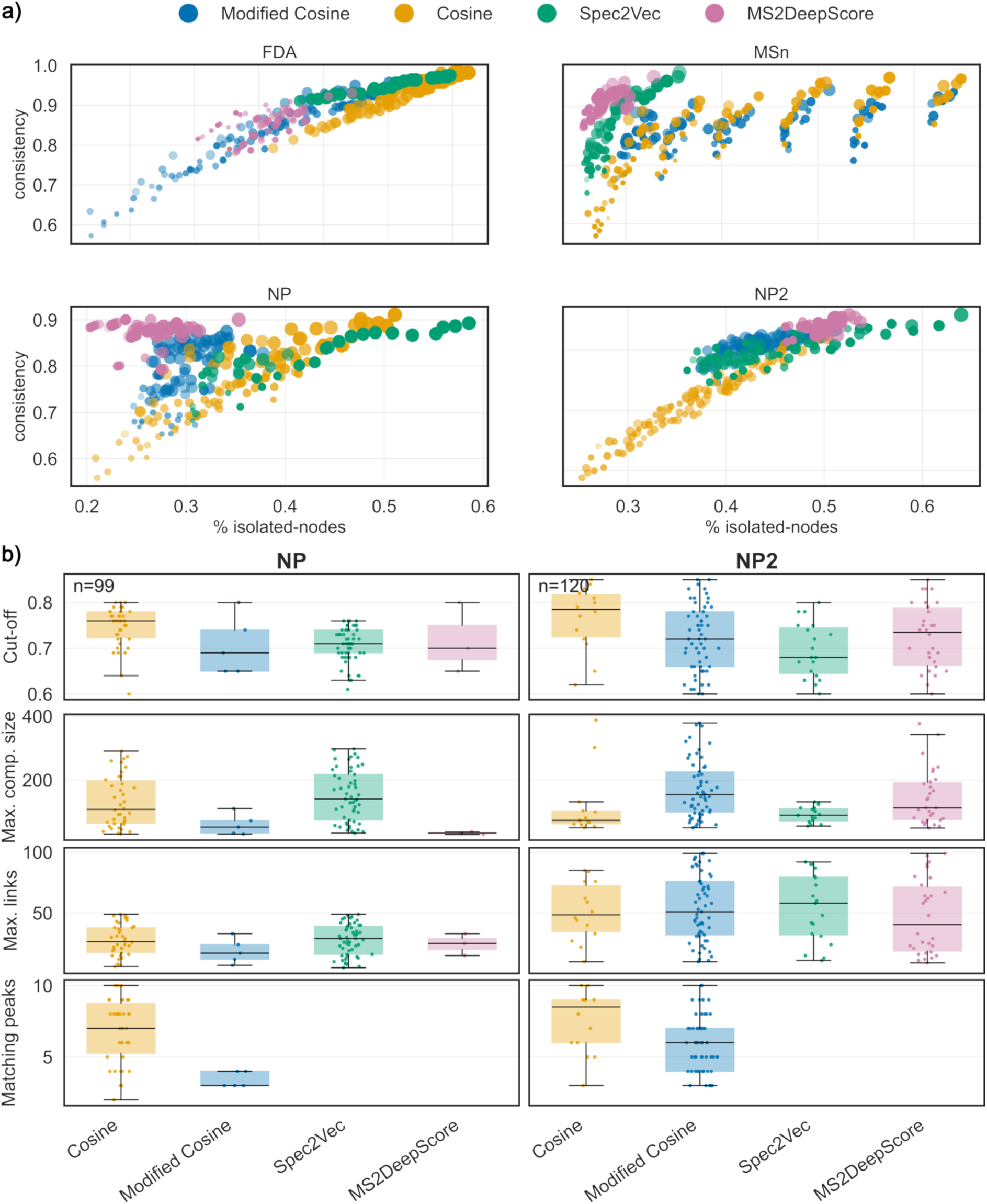
(a) Trade-off between topology and chemistry metrics. The optimal region sits on the top-left where we achieve high chemical consistency at lower percentage of isolated nodes. Dot transparency indicates CV_degree and size indicate intra-similarity. (b) Boxplots of the networks passing the thresholds: intra-similarity > 0.4, consistency > 0.8, CV of degree < 0.7, Gini < 0.5, isolated nodes < 0.55 for NP and NP2 datasets. For Spec2Vec and MS2DeepScore minimum matching peaks is not explored in this version.

If we apply one fixed target (intra-similarity > 0.4, consistency > 0.8, CV of degree < 0.7, Gini < 0.5, % if isolated nodes < 0.55) to NP and NP2, both natural-product libraries that share their dominant chemical classes (Shikimates and Phenylpropanoids, Supplementary Figure 7 and 8) but differ roughly two-fold in size, comparable fractions of networks pass all these metric thresholds in each (25% for NP, 30% for NP2). The feasible regions, however, are not interchangeable: which parameter values survive depends on the score, and for some parameters on the dataset (Figure 2b). Matching peaks and cut-off are score-driven and stable across datasets; the median surviving cut-off is highest for Cosine (∼0.8 vs ∼0.7 for Modified Cosine) and surviving matching peaks cluster around 8 for Cosine but 5 for Modified Cosine in both libraries. Maximum links, by contrast, is dataset-driven and consistent across scores, rising from a median of ∼25 in NP to ∼50 in NP2. Maximum component size shows no clear pattern, varying with both score and dataset. Hence, a configuration tuned on one similarity score would therefore fail on another similarity score even when the settings would generally look plausible, so no single parameter set is optimal across datasets or scores. This observation was precisely our motivation to develop arteMIS: i.e., to sample and rank parameters per dataset, rather than fixing them in advance with the aim to produce more useful networks with higher chemistry consistency without over-fragmenting the network.

### 2. Network optimisation and customisable ranking with arteMIS

To translate the sensitivity and feasibility analyses into actionable parameter recommendations, we ran arteMIS in global-mode, the default configuration in which every feature has a chemical annotation available and are used for the evaluation procedure. As the four benchmark datasets are spectral libraries, SMILES are already provided; therefore, MS2Query was only used to retrieve NPClassifier pathway level for the consistency metric. We combined the five-evaluation metrics into a single composite Z-score. The composite score is oriented so that higher values indicate more desirable networks: the three-topology metrics (CV of node degree, Gini coefficient, and % isolated nodes) are sign-inverted before summation, while intra-similarity and chemical consistency enter with their intrinsic positive sign (see Methods section Evaluation metrics – Selection of representative configurations).

For each dataset per scoring method combination, the 100 sampled networks were ranked by this score, and the top-ranked three configurations were used for downstream analysis. In Table 2, the highest top configuration is shown per score and dataset.

**Table 2.** arteMIS resulting top-ranked configurations by composite-score for each spectral similarity score, showing topology and chemistry-based metrics across the four evaluated datasets. Matching peaks is not applicable in this version for machine learning-based scores.

| Dataset | Max. Comp Size | Max. Links | Cut off | Matching peaks | CV degree | % Isolated Nodes | Gini-coefficient | Intra-similarity | Consistency | Similarity score | Composite Score |
| --- | --- | --- | --- | --- | --- | --- | --- | --- | --- | --- | --- |
| FDA | 82 | 10 | 0.58 | 8 | 0.75 | 0.88 | 0.06 | 0.58 | 0.97 | Cosine | 1.98 |
| FDA | 17 | 13 | 0.78 | 4 | 0.65 | 0.67 | 0.19 | 0.40 | 0.89 | Modified Cosine | 4.09 |
| FDA | 11 | 14 | 0.68 | - | 0.64 | 0.66 | 0.19 | 0.34 | 0.89 | MS2DeepScore | 5.73 |
| FDA | 78 | 7 | 0.72 | - | 0.73 | 0.84 | 0.09 | 0.53 | 0.96 | Spec2Vec | 1.26 |
| NP | 68 | 6 | 0.74 | 9 | 0.58 | 0.45 | 0.40 | 0.54 | 0.88 | Cosine | 3.44 |
| NP | 87 | 9 | 0.64 | 5 | 0.53 | 0.30 | 0.51 | 0.50 | 0.85 | Modified Cosine | 4.59 |
| NP | 49 | 6 | 0.67 | - | 0.51 | 0.27 | 0.53 | 0.50 | 0.88 | MS2DeepScore | 3.79 |
| NP | 49 | 6 | 0.67 | - | 0.56 | 0.40 | 0.44 | 0.46 | 0.82 | Spec2Vec | 2.82 |
| NP2 | 60 | 62 | 0.85 | 8 | 0.65 | 0.50 | 0.33 | 0.53 | 0.91 | Cosine | 5.16 |
| NP2 | 92 | 11 | 0.72 | 7 | 0.60 | 0.43 | 0.39 | 0.55 | 0.91 | Modified Cosine | 3.17 |
| NP2 | 56 | 64 | 0.73 | - | 0.66 | 0.53 | 0.31 | 0.58 | 0.95 | MS2DeepScore | 4.76 |
| NP2 | 50 | 29 | 0.85 | - | 0.71 | 0.69 | 0.24 | 0.60 | 0.95 | Spec2Vec | 5.6 |
| MSn | 56 | 46 | 0.79 | 5 | 0.50 | 0.05 | 0.45 | 0.57 | 0.89 | Cosine | 6.5 |
| MSn | 51 | 33 | 0.74 | 5 | 0.47 | 0.05 | 0.47 | 0.59 | 0.89 | Modified Cosine | 8.05 |
| MSn | 81 | 99 | 0.83 | - | 0.47 | 0.04 | 0.45 | 0.65 | 0.93 | MS2DeepScore | 6.02 |
| MSn | 50 | 29 | 0.85 | - | 0.47 | 0.07 | 0.40 | 0.79 | 0.96 | Spec2Vec | 5.28 |

The design choice is what makes the arteMIS ranker adjustable: researchers can re-weight or drop metrics to steer the ranking toward their specific goal. For example, increasing the weight of certain chemistry metrics (e.g. intra-similarity) if finer relationships need to be considered or by giving more importance to topology metrics when working with over-sparse networks where recovering connectivity is the priority. The unweighted composite score reported here is therefore a starting point, not a fixed recommendation.

The top composite score varies substantially across score and dataset combinations, from 1.26 (Spec2Vec on FDA) to 8.05 (Modified Cosine on MSn). By construction, the composite is a sum of Z-scored metrics, median values sit near zero (-0.24 to 0.50 across combinations). Minimal Z-values are consistently large and negative (-2.22 to-6.68), confirming that the sampling truly explores poor regions of parameter space and favorable ones. Reported minimal, medians, and maximum values are reported in Supplementary Table 2. Higher top values indicate score–dataset combinations where topology and chemistry objectives are jointly satisfiable, while lower ones indicate combinations where no sampled configuration simultaneously optimises both.

A notable trend across Table 2 is that the top composite scores tend to increase from FDA (1.26–5.73) through NP (2.82–4.59) and NP2 (3.17–5.60) to MSn (5.28–8.05), however combination medians remain close to zero across all four datasets, and minimal values are actually most negative on MSn (down to-6.68), so the pattern reflects a higher limit on larger datasets rather than a uniform increase. Therefore, joint chemistry–topology objectives being easier to satisfy on larger, where a wider spread of pairwise similarities gives the sampler more room to find configurations that produce well-connected components without collapsing the network. The configurations in Table 2 serve as the starting point for the robustness analysis in the next section.

### 3. Robustness of top-ranked molecular networks under subsampling

Mass feature detection in metabolomics is subject to run-to-run variation, MS/MS precursor selection during data acquisition is stochastic, and minor differences in sample composition can shift which features are fragmented during metabolomics experiments. A useful network configuration should therefore be robust to this biological and technical variation^33,34^.

From the configurations identified in Section 2, we selected the top three ranked configurations per spectral similarity score and dataset (12 configurations per dataset, 48 in total), and generated bootstrap network replicates (B = 1000) for each. The subsampling process works as follows: we randomly kept 85% of the spectra of each sample and, using the parameters selected, we rebuild the network. From these replicates we compute the following robustness metrics: probability of a node being isolated (the tendency of nodes to become singletons under resampling), and edge-level stability (how consistently edges observed in the original network are reproduced across bootstrap replicates, see Methods Table 7). We compared the robustness metrics between the top-configurations versus the GNPS default networks computed once (score cut-off = 0.7, minimum 6 matching peaks, maximum component size = 100, maximum links per node = 10)^9^. Reported values for top configurations are means and standard deviations across the three parameter sets, to illustrate both subsampling variation and small differences between the three top-configurations. The default configuration is a single parameter set, so it was computed once per dataset, and no standard deviation is reported for it.

For Modified Cosine, in the smallest dataset (FDA, 606 nodes), the top configurations yielded a mean node isolation probability of 0.68 (standard deviation (SD) ±0.020), compared with 0.71 of the default configuration. For the largest dataset (MSn-COCONUT, ∼13K nodes), node isolation probability was low across all configurations, with values of 0.099 (SD ±0.058), for the top-ranked, and 0.08 for the default configuration (Figure 3 (a)). This observation indicates that for larger networks the probability of node isolation is lower regardless of the parameter of choice, since in larger networks, nodes have a higher probability to be connected, increasing the overall robustness of the resulting molecular families.

**Figure 3.**
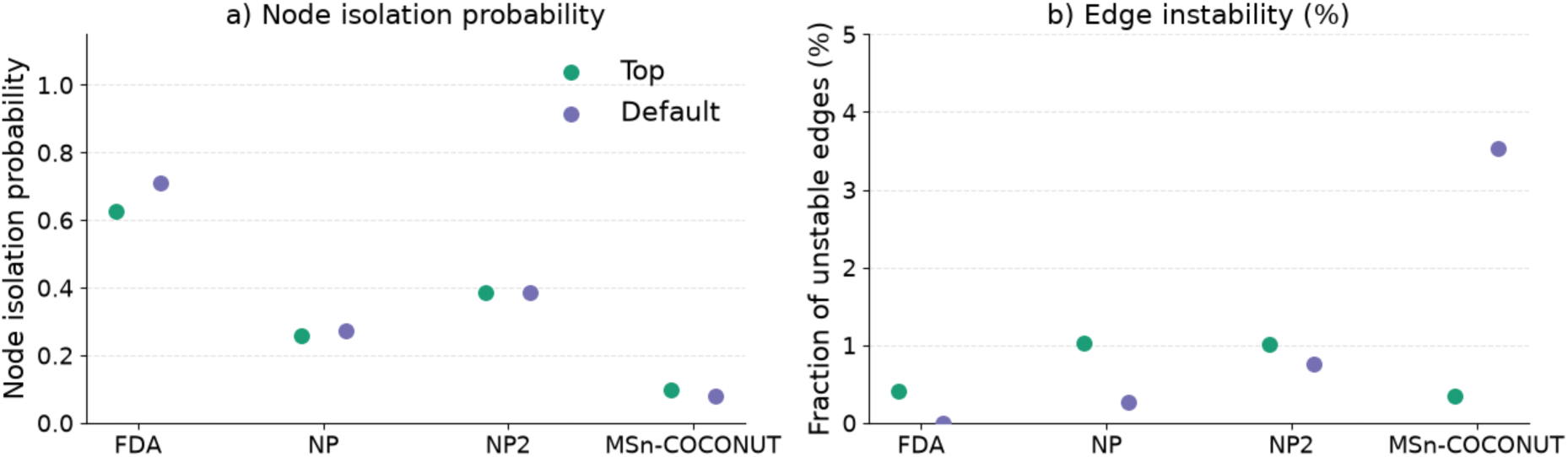
Robustness of Modified Cosine molecular networks under 85% of nodes subsampling. a) node isolation probability of top (green) vs. default (purple) configurations across datasets (FDA-dataset containing 606 spectra, NP-dataset containing 1226 spectra, NP2-dataset containing 7250 spectra and MSN-COCONUT containing 12929 spectra). b) fraction of unstable edges across datasets tops vs. defaults configurations across datasets.

Edge stability was strongly dataset-size dependent (Figure 3 (b)). In small datasets, all subsampled configurations reached a stability value of 1.0, because the networks never became large enough to cause molecular families to fragment, the maximum component size and maximum links thresholds were never reached. In larger datasets, however, the default parameters (maximum component size = 100, maximum links = 10) introduced substantial edge instability. For NP and NP2 top configuration yielded slightly higher fraction of unstable edges, but the values still less than or equal to 1% of the total edges. For the largest MSn dataset, 3.5% of original edges were inconsistently present across replicates under the default configuration, compared with 0.3% (SD ± 0.002), under the top configuration.

#### 3.1 Robustness across spectral similarity scores

For cosine, in the smallest dataset the probability of node isolation for the top configuration is the same as the default (0.88 SD ± 0.013). For the MSn-COCONUT largest dataset, the probability of a node being isolated is also lower, top giving a 0.1 (SD ± 0.060) and default 0.08. For the edge stability again showed a size-dependent behaviour, no effect in smaller datasets but in the largest datasets 1.9% of the edges were not reproducible under resampling for the default configuration but in the top the inconsistency is reduced to 0.2% (SD ± 0.003) (Supplementary Table 3).

For Spec2Vec, for the smallest dataset the probability of node isolation is also a bit higher for the top configuration (0.85 vs 0.84). For the largest dataset (MSn-COCONUT), the edge instability remained <1% for both configurations. MS2DeepScore closely followed the trend observed for the probability of node isolation in the smallest dataset (top 0.56 SD ± 0.020 vs 0.48 from default). In this case, default generates of 2.0% unstable edges, while 0.1% (SD ± 0.001) of the top configurations (Supplementary Table 3).

Overall, as noticed, top and default configurations generally produced similar results for probability of node isolation across datasets and scores. The the largest differences observed where regarding edge instability, on MSn-COCONUT, which was roughly 10x lower in the top-configurations networks compared to defaults across scores.

### 4. Case Studies: Focused Optimisation Modes

The two case studies below showcase two of arteMIS’ annotation-guided optimisation modes. In seed-mode, chemistry metrics are computed only over the subset of features with reliable annotations, so the parameter search is guided by the chemical information of the annotated features of the network while the rest of the topology keeps being the same. In target-class mode, optimisation is driven specifically by the co-clustering of a chosen chemical class. The two modes address different questions:’give me the best network for the compounds I already know’ versus’give me the best network for a certain class of compounds’, and, as we show, converge on different parameter regions even on the same dataset.

#### 4.1 Seed-mode application

To showcase the utility of arteMIS seed-mode in a real-world scenario, we applied it to a new dataset of 101 actinobacteria strains spanning six genera (*Streptomyces, Microbispora, Actinoplanes, Dactylosporangium, Planomonospora*, and *Micromonospora*), comprising 10,326 features in total. Of these, 166 were structurally annotated (for details check Methods – Case Study seed-mode) and served as our ground truth; pathway-level classifications of these compounds were retrieved from NPClassifier. These annotated features served as “seed” to optimise network parameters and evaluate topology and chemistry metrics. In seed-mode, arteMIS calculates chemistry metrics exclusively from the well-annotated features, directing optimisation toward the molecules of interest, rather than tuning parameters globally across a largely unlabelled dataset. The primary aim is to obtain molecular families that are more chemically meaningful for the annotated compounds compared than those obtained with default network settings.

#### Network selection based on arteMIS composite-score

Table 3 compares the default network with the top-three configurations identified by arteMIS. Among these, top1 was selected for further analysis as it achieved the highest composite score (4.15); although top2 scored similarly, top1 was preferred for its lower percentage of isolated nodes. Compared to the default, top1 outperformed across all topology and chemistry metrics: Specifically, the percentage of isolated nodes decreased from 0.43 to 0.38, while intra-similarity increased from 0.63 to 0.67. These results demonstrate that arteMIS can balance the usual topology-chemistry trade-off to achieve improvements on both sides.

**Table 3.** Default network (standard settings) and top-ranked networks identified by arteMIS in seed-mode.

| Rank | Max. comp size | Max. Links | Cut-off | Matching peaks | CV_degree | % isolated nodes | Gini-coefficient | Intra-similarity | Consistency | Composite score |
| --- | --- | --- | --- | --- | --- | --- | --- | --- | --- | --- |
| 1 | 131 | 10 | 0.57 | 8 | 0.60 | 0.38 | 0.46 | 0.67 | 0.99 | 4.15 |
| 2 | 71 | 6 | 0.78 | 9 | 0.60 | 0.47 | 0.39 | 0.67 | 0.99 | 3.92 |
| 3 | 142 | 9 | 0.7 | 8 | 0.61 | 0.44 | 0.41 | 0.67 | 0.99 | 3.41 |
| default | 100 | 10 | 0.7 | 6 | 0.61 | 0.43 | 0.41 | 0.63 | 0.99 | - |

Moreover, at the global level (Figure 4), top1 had 13,672 edges versus 10,989 than the default, and reduced singletons from 4,095 to 3,580; the network becomes denser and less fragmented under the optimal arteMIS settings.

**Figure 4.**
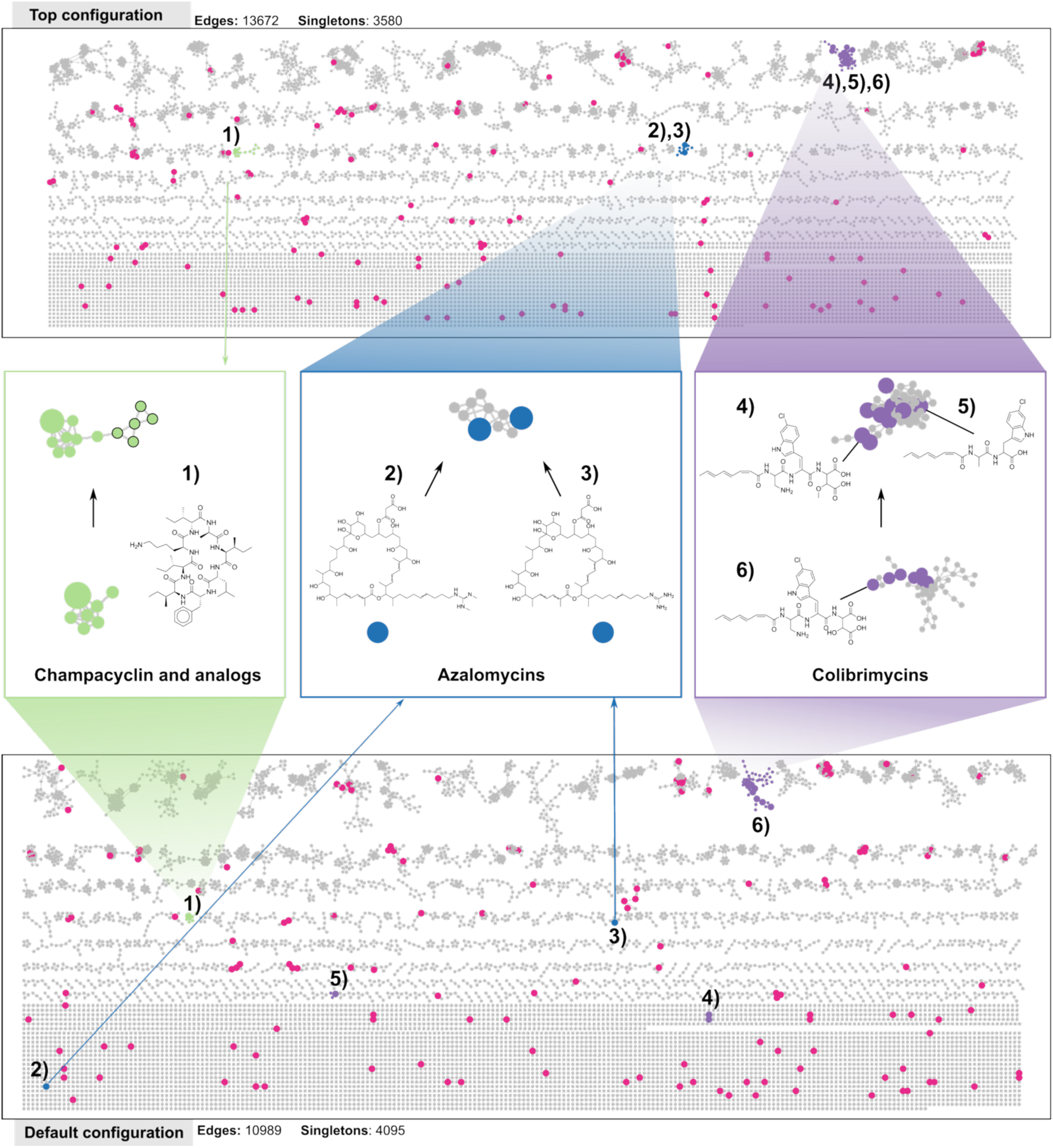
Case Study arteMIS seed-mode using Modified Cosine comparison top1 vs GNPS-default settings (bottom). On top the top1 configuration network found by arteMIS by optimizing the five metrics: i.e., reducing three topology (coefficient of variation of node degree, gini-coefficient and % of isolated nodes) and maximising two chemistry-based metrics (intra-similarity and chemical NPClassifier-consistency). On the bottom, the default network consisting of 10,989 total edges and 4,095 singletons while the top-ranked network increased the number of edges to 13,672 and reduced the number of singletons to 3,580. In the middle row, we highlight three example molecular families that were improved: the champacyclin-family (left) that increased from 7 to 13 family members (in bold) potentially annotated as “oligopeptides” (in green); the azalomycin-family (middle), that reunited azalomycin 5fa and f3a singletons into a single family in top1, and the colibrimycin-family (right) that reunited different analogues that previously were fragmented in multiple families under the default settings.

#### Changes in molecular family composition

At the molecular family level, we first tracked how the clustering of the 166 annotated compounds changed between the default and top1 networks: 43 annotated nodes now belonged to a larger family, 33 to a smaller one, and 90 remained unchanged (see Supplementary Table 4 for full composition details).

One of the most common outcomes during network construction is the singleton node generation, that can be caused by parameter choices (for example high similarity cut-off), or by the nature of the data used (no other structurally similar molecules). Singletons cannot propagate structural information to neighbours, one of the key uses of molecular networking, so rescuing them is important for downstream annotation propagation.

Notably, from the 55 annotated features that were singletons in the default network, 9 structurally different nodes were successfully rescued in top1, including azalomycin, nigericin, pamamycin-691, different acyl-deferroxamines, terragine and 1-1-hydroxy-6-methoxy-phenazine (Supplementary Figure 9). For example, in the top1 configuration, azalomycin F5a (2) occurs in the same molecular family as azalomycin F3a (3) (Figure 4). In the default network, the edge between azalomycins did not occur because of the cut-off of 0.70; with top1 lower cut-off (0.57) arteMIS captured a spectral similarity of 0.60 between fragmented molecules generated an edge and connected these molecules in the same molecular family. This shows that fine-tuned selection of parameters can be more effective than default settings for recovering members of specific metabolite family and it also shows that reducing the percentage of isolated nodes can result in bonafide edges in the molecular network rather than just false positive links.

Beyond singleton rescue, several families already connected in the default network increased in membership with structurally related molecules in top1, including colibrimycin, nigericin and chrymostatinol. The colibrimycin family, for instance, expanded from 7 annotated nodes in the default to 11 in top1. Figure 4. shows in the default network, some of the adducts (colibrimycin C6 [M+NH^4^]+, colibrimycin A1 [2M+H]+ and colibrimycin A1 demethyl [M+H]+; 4), 5) and 6)) were fragmented among different molecular families. Here, cut-off is not the cause of the separation (edges show >0.9 similarity), and neither is the maximum component size cap, since the family size in both families is ∼50 members and maximum links (is equal to 10 in default and top1). The most plausible explanation is that during the 100-network construction, this combination of parameters produced higher intra-similarity than the default, which is what pushed it to the top.

Beyond improving the grouping of known compounds, arteMIS seed-mode can improve the connecting with unknown but potential mass features. For instance, in Figure 4, champacyclin (node ID: 6557, 1)), a cyclic octapeptide, went from a 7-member family in the default to a 13-member family in top1. We do not have manually annotated compounds as neighbours here, but the surrounding nodes were annotated by MS2Query as “oligopeptides” (all with >0.7 analogue similarity), with most features corresponding to surugamide G and its analogues (A, H, and others). Further analysis revealed that champacyclin and the surugamides share a cyclised octapeptide scaffold (Supplementary Figure 10). This showcases that arteMIS not only improves the regrouping the known molecules but also improves the grouping with potentially similar molecules.

We acknowledge that not all molecular families did improve; we found two specific cases, marinomycin A and TPU-0037-B, where in the top configuration these compounds were fragmented from their previously coherent families. The two cases arose for different reasons.

Marinomycin A (node ID: 7366, adduct [M+H]+) belonged to a structurally coherent family in the default network, including another annotated marinomycin A of a different adduct ([M-H₂O+H]⁺, node ID: 7365). The edge linking 7366 to this component shared seven matching peaks (Supplementary Figure 11), enough to pass the default minimum of six, but not top1’s minimum of eight. Under top1, that edge is filtered out and the node is left isolated. One could argue for simply lowering the matching-peaks threshold, but a lower threshold can just as easily let structurally unrelated molecules into a family. A concrete example: in the default network, piericidin A and sesbanimide A sit in the same larger family, yet piericidin A is a flat pyridine ring^35^ attached to a long unsaturated hydrocarbon tail while sesbanimide A is a saturated glutarimide linked to a small oxygen-rich ring system^36,37^. They are also produced by different polyketide machineries (piericidin by a cis-AT PKS, sesbanimide by a trans-AT PKS^38^), so grouping them together is not supported structurally or biosynthetically. Under top1 they sit in two separate families, each with more plausible neighbours. In short, the stricter matching-peaks threshold can be useful elsewhere in the network, keeping unrelated nodes separated.

TPU-0037-B fragmentation (node IDs 5962 and 5637) is a more interesting failure, and worth mentioning because it points to a parameter that we currently under-explore. The edge between the two isomers had a cosine of 0.851 and shared 18 matched peaks: it survives the score cut-off, the peak-matching filter, and the maximum links cap. However, it got removed by the maximum component size step. When a family exceeds the maximum component size, arteMIS raises the score threshold in fixed increments of cosine_delta (0.05), starting from cut-off, until the component is small enough. Because the default and top1 begin from different starting points (0.70 vs 0.57), the ladders sit on different final thresholds, 0.85 for the default (which keeps the 0.851 edge) and 0.87 for top1 (which cuts it). The edge is lost not because it has a low similarity score but because 0.85 is simply not a reachable step when the ladder starts at 0.57. This is a discretisation artefact rather than a chemistry failure, and it identifies cosine_delta as a parameter worth including in future arteMIS versions.

This seed-mode case study shows what arteMIS can achieve in practice. The systematic parameter exploration revealed that even with a relatively relaxed spectral similarity cut-off of 0.57, the top1 network outperformed the default network across all evaluated topology and chemistry metrics. The optimized settings rescued at least 9 annotated nodes from singleton status, expanded several existing molecular families with structurally related members, and fragmented only two families: one due to a stricter matching-peaks filter (which appears useful elsewhere in the molecular network), and one due to a discretisation artefact in cosine_delta that points to a clear next step for arteMIS development. By operating in seed-mode and focusing optimisation on the subset of chemically annotated molecules, arteMIS directly improves clustering of known compounds and their close analogues present in the dataset. In practice, this means that with a modest set of standards and a large pool of unannotated features, one can optimise the settings to better group these features and then propagate the structural information from the standards to plausible neighbouring nodes.

#### 4.2 Target class-mode application (melleolides)

To further illustrate the capabilities of arteMIS, we implemented an alternative usage of the optimisation framework for targeted chemical class tuning, in which the hyperparameter optimisation via Latin Hypercube Sampling is guided not by global network chemical evaluation metrics, but by the clustering behaviour of a specific compound class of interest (target-class mode).

For this purpose, we revisited the metabolome of *Armillaria ostoyae*, a prolific fungal producer of the bioactive meroterpenoids known as melleolides. In our previous pilot study^39^, melleolide chemical diversity was captured by propagating annotations of isolated metabolites within the FBMN. Despite the high structural similarity shared across all melleolide analogs that feature a sesquiterpene protoilludene backbone esterified with an orsellinic acid moiety^39–41^, these compounds were distributed across distinct molecular families, highlighting the fragmentation problem that targeted chemical class mode is designed to address: is it possible to reunite analogue features otherwise scattered across the molecular network?

The above issue motivated us to evaluate whether arteMIS could reconcile melleolide diversity into fewer, more coherent molecular families. We used the annotated melleolides to drive the network optimisation (including singletons), with the goal of identifying network parameters that maximally co-cluster melleolide-like nodes. This approach builds on an ongoing assumption that we use in global mode: for a metabolome, few metabolites are true (singletons) islands, while most compounds are part of biosynthetic pathways or arise during chemical transformations of related molecules. Therefore, minimising singletons is a biologically motivated objective during molecular network construction.

As observed in Figure 5, the network generated with default settings originally had 1,031 edges, while the arteMIS optimized network (Figure 5, bottom) increased the number of edges to 1,401. After target-class optimisation, the largest molecular family included 171 nodes including most structurally similar monomeric melleolides featuring their characteristic 6-protoilludene backbone with diverse orsellinic acid substitutions as expected, as shown for melleolide B and J (Figure 5, structures (1) and (2), respectively). The target-class optimisation provided a substantial improvement compared to the default scenario, where these molecules were scattered across 24 different molecular families and the top1 network rescued 10-members into the largest family. Other types of melleolides, such bismelleolide EH (3) and 10-hydroxy-5’-O-methylarmillane (4), remained in a molecular family comparable to the one observed in the default network. From a total of 20 melleolide nodes that were singletons in the default network we were able to rescue 10 nodes to molecular families that were composed of similar molecules.

**Figure 5.**
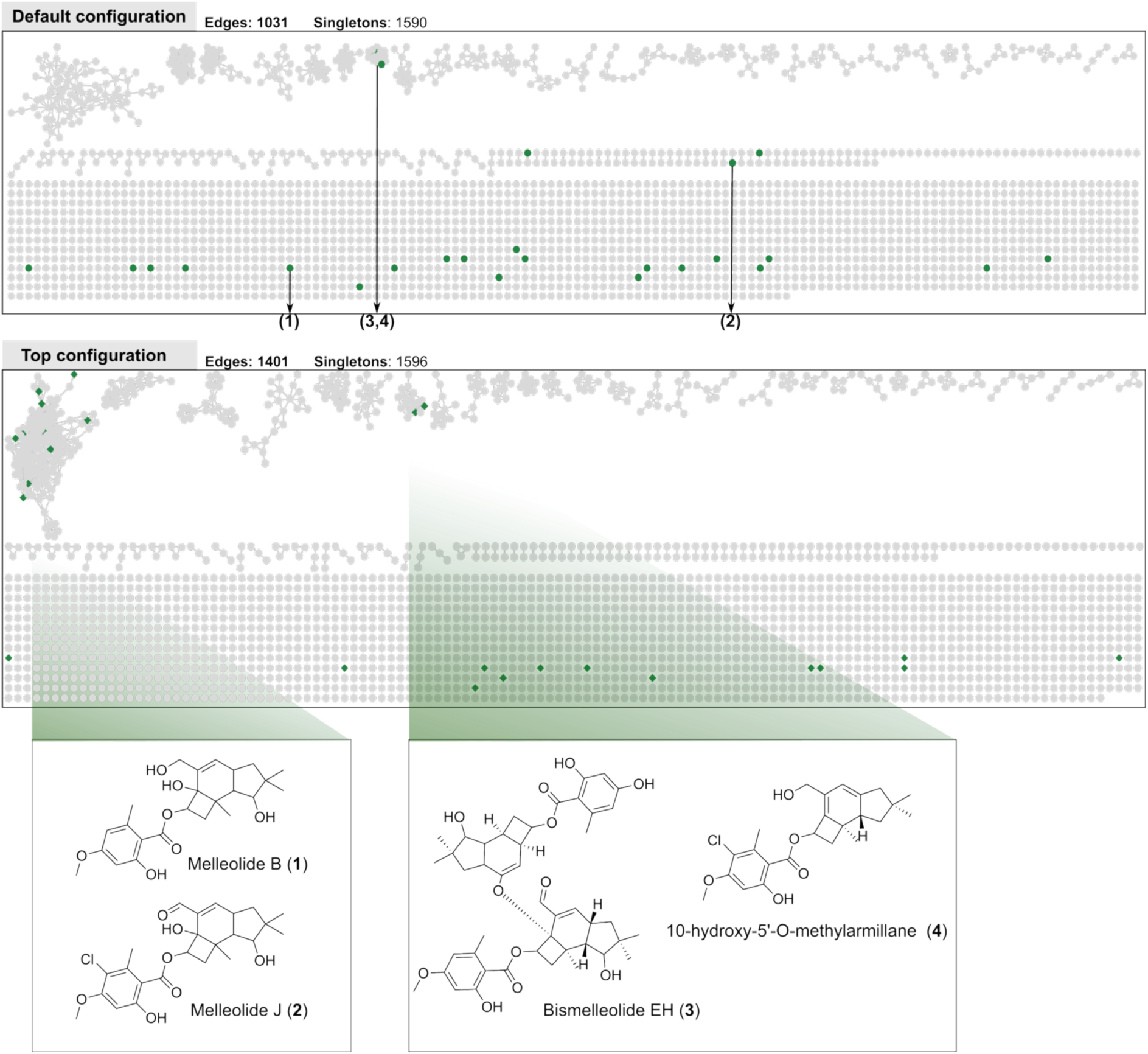
Case Study on melleolide targeting class-mode. As in the original study^39^, using the default settings, the melleolide-like molecules were fragmented into 24 molecular families, where only 5 molecules were not singletons. arteMIS generated a top-ranked network where, we were able to connect these singletons back to the largest-family composed of 171 nodes, joining back 10 previous melleolide-like molecules that were singletons.

We also compared the results of the target-mode versus the global-mode. The global-mode top1 network rescued one more singleton melleolide compared to the default network (see Supplementary Figure 12), but the gains obtained in target-mode were superior. Beyond the numerical improvement, the two modes converged in different parameter combinations, notably with a cut-off of 0.8 versus 0.62 (for further details see the parameter top1 from global-mode in Supplementary Table 5), global-mode has to optimise mass features and thus may compromise performance for specific classes. These results have broader methodological implications, as the target-class mode demonstrates that network parameters governing molecular family construction are not universally optimal but are instead inherently dependent on the structural diversity and characteristic MS/MS patterns for distinct metabolite classes^26^. By allowing users to tailor this optimisation framework to a chemically meaningful set of annotations, arteMIS effectively reframes parameter selection from an unsupervised global problem into a supervised, class-aware one.

The above addresses a longstanding challenge in untargeted metabolomics, where the high number of unknowns has made it difficult to conduct analyses focused on targeted discovery of specific natural products of interest. In contrast to genome mining, where biosynthetic gene clusters provide a natural and explicit scaffold for targeted compound discovery^40,41^, metabolomics workflows have traditionally lacked analogous mechanisms for directing network analyses toward compounds of interest. We expect that arteMIS contributes to closing this gap by enabling various analysis modes: global, seed and target-class that are simultaneously data-driven and hypothesis-guided. Rather than replacing expert knowledge, the targeted chemical class mode facilitates it as existing annotations are employed as anchors to recover and organide related but previously disconnected metabolites.

We anticipate that the target-class mode will prove particularly valuable in discovery pipelines, where the exploration of chemical diversity is often challenged by the complexity of natural products and the limited availability of reference spectra. More broadly, we expect that class-aware network optimisation frameworks such as the here introduced arteMIS in target-class mode will strengthen the integration of metabolomics with large-scale genome mining campaigns^42,43^, where the ability to connect biosynthetic gene clusters of interest across complex metabolomes remains a critical bottleneck^44^.

## Conclusions and Outlook

The arteMIS framework introduced here provides a principled, customisable framework for molecular network construction that replaces default-parameter reuse with goal-directed optimisation. Across four datasets and four scoring methods, arteMIS-ranked configurations matched or outperformed GNPS defaults in chemistry-based, topology-metrics and in edge-level robustness under subsampling, with the largest gains on datasets of more than ten-thousand spectra. The two case studies illustrated concrete benefits in both seed-mode (rescuing nine annotated singletons and expanding structurally meaningful families in an actinobacterial dataset) and target class-mode (rejoining fragmented melleolide analogues in a fungal species), complementing the global-mode benchmarked used across the four library datasets.

This framework provides three complementary modes made for different experimental starting points. Global mode is applied as default, and it is used when every feature has chemical annotations, either provided directly (as in curated spectral libraries) or predicted through MS2Query; it was used for benchmarking (using spectral libraries), method development and datasets where a broad coverage exists. Seed mode is intended for exploratory datasets with a curated subset of high-confidence annotations, the goal is to propagate structural context from these anchors (such as standards). Target-class mode prioritises the recovery of a specific compound class, and is most useful when the biological question is class-focused, such as tracking a specific class of natural products through a producer organism. The main differences between seed and target-class modes are that target-class weights class recovery, while seed modes weights connectivity of the annotated anchors independently of their class. Beyond the tool itself, our benchmark establishes a methodological point that we believe has been underappreciated so far: for molecular networking, parameter selection is as important as the choice of similarity score, and optimal settings do not transfer between scoring methods or dataset sizes.

We acknowledge several limitations of our framework. First, no finite set of metrics fully captures all desirable network properties: topology and chemistry exist in tension, and the fingerprint-based metrics we use can overstate similarity when substructures are widely shared or understate it under stereo-or regiochemical variation^45^, and some metrics are limited in their capacity to cover topological and chemical properties. For instance, Adjusted Mutual Information (AMI) used to measure partitioning alignment, penalizes chemical class partitioning, however, chemical classes are shared across different molecular families (for further explanation see Methods – Evaluation metrics). Another one is gini-coefficient, which prevents hairball formation (reduces the inequality of molecular families), however a perfect gini-score (equal to 0, all molecular families being modular) could simply imply highly fragmented molecular families or all nodes acting as singletons. In practice, users can mitigate this by pairing the composite score with manual inspection of a small number of the top-ranked configurations, as we have shown in the case studies.

Second, the composite Z-score depends on the choice and weighting of metrics, which users should align with their biological question; the unweighted form we report here is a neutral default, not a universal recommendation. Users should align weights with their biological question. For instance, on an analog library dataset, in global-mode, intra-similarity could be up-weighted relative to chemical consistency (NPClassifier), since most molecules are expected to share structural cores. In contrast, on datasets composed of unrelated molecules, chemistry metrics could penalize the network topology, and increasing the weights of the topology metrics gives arteMIS more flexibility to connect distant nodes.

Third, the parameter space explored by arteMIS is not fully symmetric across scores: minimum matching peaks is optimised for the cosine-based scores but not for MS2DeepScore or Spec2Vec. Therefore, configurations for the ML-based scores are ranked over a slightly smaller parameter space (see Methods). This asymmetry is not a design choice but reflects the current state of the underlying libraries, since in matchms^46^, and this can be added symmetrically once the information becomes accessible. Finally, the maximum component size step currently raises the score threshold in fixed increments of cosine_delta (Δ = 0.05) anchored to the user-defined cut-off, so two configurations starting from different cut-offs land on non-overlapping ladders (as illustrated by TPU-0037-B in Section 4.1); refining this increment, anchoring it to an absolute grid, or including cosine_delta in the LHS sampling would remove the artefact.

Looking forward, arteMIS could be extended in several directions. First, mapping unstable edges and isolation-prone nodes back onto the network topology, and linking those regions to chemical annotations or spectral quality (such as low fragment counts, where each peak carries more weight in the similarity scores), could reveal whether particular compound or spectrum types are systematically more susceptible to run-to-run variation. Second, integrating community-detection approaches that move beyond fixed-parameter pruning, a recent application to metabolomics has shown that identifying and pruning natural molecular communities can raise connectivity from ∼25% to ∼95% of the features; incorporating community-detection methods in our framework could be a next step^47^. Third, molecular-family coherence could be limited based on exact mass differences, where users could define a maximum tolerated mass difference within a molecular family (e.g., 180 Da), hypothesizing that molecules with substantially divergent exact masses are less likely to belong to the same coherent chemical family, mass-difference networking has a long history in metabolomics^48^ and biotransformation-consistent edge annotations have recently been applied at scale to plant metabolomes^49,50^, suggesting a concrete integration path. Fourth, edge-level confidence assessment could be coupled to arteMIS in a two-stage pipeline in which arteMIS first optimises global parameters and a tool such as SpecReBoot^51^ then quantifies the reproducibility of individual edges within the resulting network.

In general, arteMIS is designed to be modular and customisable. The framework is built on top of matchms and produces networks that can be exported directly to Cytoscape, allowing users to incorporate arteMIS into existing untargeted metabolomic workflows (incorporates multiple scores) between spectra processing and downstream analysis. Therefore, arteMIS-optimised networks can be used for annotation propagation, substructure discovery (Mass2Motifs) mapping, bioactivity mapping, and the linking of biosynthetic gene clusters to their metabolomic counterparts in genome-mining campaigns, a current bottleneck in the natural products field. Together, the global, seed and target-class mode make arteMIS applicable across a spectrum of experimental settings, from fully unannotated exploratory datasets to class-focused discovery, while keeping its metrics customisable and their weighting open to the user. Here demonstrated with natural products benchmark datasets and case studies, we expect that our arteMIS framework is applicable across different metabolomics disciplines including clinical and environmental metabolomics studies. We anticipate that treating parameter selection as a data-driven, hypothesis-optimisation problem, rather than a fixed parameter selection, will make molecular networking outputs more useful, more reproducible, and more reliable across the untargeted metabolomics community.

## Methodology

### Datasets, preprocessing and scores

All spectral preprocessing (benchmark library datasets and case studies) was performed using matchms^46^ for cosine-based metrics that include matching peaks as input, we used the branch development (https://github.com/matchms/matchms/tree/development), and for the machine-learning similarity scores we used the standard version 0.31.0. All spectra were filtered using the default matchms filtering pipeline: SpectrumProcessor (DEFAULT_FILTERS + CLEAN_PEAKS).

The DEFAULT_FILTERS + CLEAN_PEAKS pipeline combines metadata harmonisation with spectrum-level peak cleaning. First, DEFAULT_FILTERS standardises metadata fields, filling missing fields where possible, drops spectra lacking essential metadata, and normalizes peak intensities so the base peak equals 1.0. Then CLEAN_PEAKS restricts peaks to the 0–1000 Da range, removes low-intensity noise below 0.1% of the base peak, discards peaks above the precursor m/z, caps each spectrum at the 1000 most intense peaks, and only centroid-mode spectra is kept, it also removes instrument-specific noise bands. A quality gate requires at least 5 peaks with ≥2% relative intensity, otherwise the entire spectrum is dropped.

Four different datasets of increasing size and chemical diversity were used for our analyses. The FDA-dataset consists of reference compounds generated by the Dorrestein Lab (GNPS-SELLECKCHEM-FDA-PART2.mgf). The original dataset contained 656 spectra, which were reduced to 606 spectra after matchms filtering. The NP-dataset was obtained from the NIH Natural Products Library (GNPS-NIH-NATURALPRODUCTSLIBRARY.mgf), also generated by the Dorrestein Lab. This dataset initially contained 1267 spectra, of which 1226 spectra remained after matchms filtering. The NP2-dataset corresponds to the second release of the NIH Natural Products Library (GNPS-NIH-NATURALPRODUCTSLIBRARY_ROUND2_POSITIVE.mgf). The original dataset included 7915 spectra, from which 7250 spectra remained after matchms filtering. Finally, the MSn-COCONUT dataset was constructed from a harmonized and cleaned collection of MS/MS spectra derived from GNPS and the recently available MSn-Lib^52^ (https://zenodo.org/records/16882111). The MSn-COCONUT originally had 1,017,531 total spectra. From this collection, the dataset was filtered to contain only MS2 data of positive ionisation mode. To restrict the dataset to natural products, annotated spectra were matched by InChIKey against the COCONUT^53^ database (https://zenodo.org/records/10629838), with a total of 407,029 registered compounds in the database. Only mass spectra of compounds present in COCONUT were retained, resulting in a final selection of 15,947 spectra. After matchms filtering, 12,929 spectra remained for downstream analyses.

For all four datasets, pairwise spectral similarity was computed using both Cosine and Modified Cosine similarity (tolerance = 0.02) using matchms. In addition to the cosine-based similarities, we computed learned similarity scores using Spec2Vec^28^ and MS2DeepScore^29^. Spec2Vec similarities were calculated using the models re-trained specifically for positive ionisation mode spectra using as source the cleaned positive spectra from https://zenodo.org/records/12543129, files associated can be downloaded here: https://zenodo.org/records/15857387. MS2DeepScore similarities were computed using the publicly available pre-trained model (https://zenodo.org/records/17826815). See https://github.com/matchms/ms2deepscore and https://www.biorxiv.org/content/10.1101/2024.03.25.586580v5 for more details.

### Chemical class retrieval

For evaluating the networks from the chemical side, we predicted the chemical class annotations using NPClassifier^31^, as implemented in MS2Query (1.5.4 version), calculated from the corresponding annotated SMILES for each compound (script available in repository). In NPClassifier there are three main levels: pathway, superclass and class. For all datasets the pathway level was the one used for the chemical metric calculation.

### Parameters used

Following the GNPS-based workflows, we used the most common user-controlled parameters: the similarity scores threshold (cut-off), maximum component size, maximum links and minimum matching peaks, described in Table 5. Bellow the order in which the parameters were applied to the network; for further details revise the code.

**Table 5.** Molecular network construction parameter explored and implemented in arteMIS.

| Parameter | Description | Acronym |
| --- | --- | --- |
| Similarity score threshold (cut-off) | Minimum similarity value (cosine, modified cosine, SpecVec or MS2DeepScore) between nodes required to draw an edge | Cut-off |
| Maximum component size | Maximum number of nodes allowed in molecular family; if the value is exceeded, the weakest edges are iteratively removed in steps defined by cosine_delta (for now steps of 0.05 across these experiments) | Maximum component size |
| Maximum links | Maximum number of neighbours retained per feature, keeping the strongest connections first | Maximum links |
| Minimum matching peaks | Minimum number of matching peaks between two spectra in order to draw an edge | Matching peaks |

### Quasi Parameter Sampling

To cover as much of the parameter space as possible (cut-off, maximum component size, maximum links and matching peaks), we applied Latin Hypercube Sampling to generate 100 different parameter combinations. This sample size was chosen as a balance of achieving low-discrepancy values of the parameter space and keeping the computational cost of building and evaluating 100 networks for four different scoring methods within a feasible time. Sample uniformity was assessed using the centred discrepancy (CD) computed on the unit-scaled samples with scipy.stats.qmc.discrepancy (1.16.2 version, default method = “CD”). In this implementation, the discrepancy is a positive number, with 0 indicating ideal uniform coverage (lower is better). The resulting discrepancy was 0.000531 for the cosine-based methods (which additionally samples the minimum matching peaks parameter), and 0.00023 for machine-learning methods across all datasets; this indicates that 100 samples already provide near-uniform coverage of the parameter space (close to zero). At the same time, network construction and evaluation were traced, with total run times for all 100 networks per dataset ranged from 15 minutes on the smallest dataset (∼600 spectra) to under 1.41 hours (cosine-based) and 2.3 hours (ML-based) on the largest dataset (∼13,000 spectra after cleaning), total running times are reported at Supporting Table 1.

Molecular networks were then constructed as defined by each parameter combination, resulting in a total of 100 networks per dataset and similarity score.

**Table 6.** Parameter ranges for the four-dataset benchmarking explored under this study.

| Parameter | FDA dataset | NP dataset | NP2 dataset | MSn_COCONUT |
| --- | --- | --- | --- | --- |
| Maximum component size | [10,100] | [30,300] | [50,400] | [50,500] |
| Maximum links | [5,20] | [5,50] | [8,100] | [8,100] |
| Cut-off | [0.55,0.8] | [0.6,0.8] | [0.6,0.8] | [0.65,0.85] |
| Matching peaks | [2,8] | [2,10] | [3,10] | [3,10] |

### Network construction

Pairwise similarity scores were used to construct undirected molecular networks using a custom NetworkX-based implementation (version 3.5)(SimilarityNetworkMod.py, available in our repository). Nodes correspond to spectra (identified by feature_id) and edges represent spectral similarity relationships, with the similarity score stored as the edge weight.

Parameters are applied to each node’s candidate neighbours in the following order: (1) the top N candidates per spectrum are retrieved from the similarity matrix, with N set to 50 for FDA and NP datasets and 100 for the NP2 and MSn-COCONUT datasets; making sure that the pool of candidates is never smaller than user-predefined maximum links cap; (2) three conditions are then combined in a single AND: cut-off is as the minimum similarity, the minimum matching peaks filter (for cosine based scores) and the self-links are discarded; (3) among the surviving candidates, only the strongest connections are retained, up to the maximum links cap; and (4) once the full graph is built, the maximum component size is applied by removing the weakest edges within oversize molecular families until the number of nodes is reached (for parameter description see Table 5.).

After this initial graph construction, connected nodes exceeding the maximum component (molecular family) size threshold were pruned by iteratively removing low-weight edges within oversized components until all components satisfied the size constraint. Unless otherwise specified, isolated nodes were retained in the final networks.

### Evaluation metrics

#### Topology metrics

##### Coefficient of variation of node degree

The coefficient of variation of node degree is used to quantify the homogeneity of the node-degree-distribution. Lower values indicate a more homogeneous distribution of connections across nodes, whereas higher values indicate that a small number of nodes have disproportionately many connections, consisting of hub-dominated molecular networks (hairball behaviour). To normalize the metric an interval (0 to 1) for the composite score, and limit the influence of extreme hubs, the raw coefficient is rescaled by 1+CV. It is defined as:

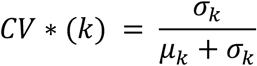

Where k_i_ is the degree of node i and:

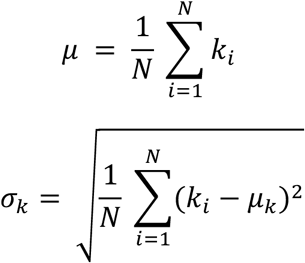

##### Fraction of isolated nodes

The faction of isolated nodes is defined as the percentage (in fraction from 0 to 1) of nodes that have zero degree, relative to the total number of nodes in the network, where 0 indicates that every node participates in at least one edge, and 1 indicates that the network does not contain connections, all features are singletons. It is defined as:

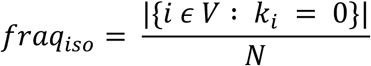

Where V is the set of nodes of network G, N = |V| is the total number of nodes, and k_i_ is the degree of node i.

##### Gini Coefficient

Inspired by the N50 metric used in genomics and adapted for molecular networking evaluation by Wang et al. (2024), we employed the Gini Coefficient, to characterise the distribution of component sizes within a network. Values close to 0 indicate that all components have a similar size, while values approaching 1 imply that one or a few molecular families dominate the network. The Gini Coefficient is computed using the following standard formula:

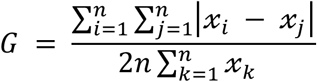

Where x*_i_* is the size of each connected component i and *n* is the total number of components, and j and k are summation indices running over all components.

#### Chemistry Metrics

##### Intra Similarity

To assess the chemical coherence within molecular families, we computed pairwise structural similarity between molecules assigned to the same network component. Molecular structures were represented using 4096-bit Morgan binary fingerprints using the RDKit^54^ toolkit (2023.9.4 version) generated from their corresponding SMILES. Pairwise Tanimoto similarity was then calculated for all molecule pairs with valid fingerprints. Molecules for which fingerprints could not be generated were excluded from this metric.

The intra-component similarity is computed as the mean Tanimoto similarity across all intra-component molecule pairs pooled across the entire network, rather than as an average of per-component means. Therefore, larger components contribute proportionally more pairs to the metric. Let C denote the set of components, M_c_ the set of molecules with valid fingerprints in component c, and T(i,j), the Tanimoto similarity between fingerprints of molecules i and j. The intra-similarity is defined then as:

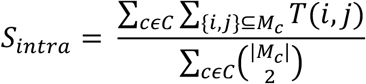

##### Consistency Measurement with NPClassifier

To evaluate the chemical meaningfulness of molecular networking components, we adopted a molecular family consistency metric inspired by Wang et al. (2024). Whereas Wang et al. used ClassyFire^55^ annotations, we relied on NPClassifier pathway-level annotations obtained via MS2Query, so the two metrics are conceptually similar but not directly comparable. This metric quantifies the proportion of nodes that belong to molecular families (components) whose nodes predominantly share the same pathway-level annotation.

For each network component *C* that contains *n* nodes, the purity P(C) is defined as the fraction of nodes belonging to the most frequent NPClassifier category within that component:

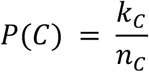

where *k_C_* is the number of nodes assigned to the most frequent NPClassifier (pathway-level) annotation.

A component is considered correctly classified if its purity meets or exceeds a predefined threshold (*τ*), set to 0.7, following work by Wang et al. (2024). This threshold tolerates minor misclassifications and allows the inclusion of related analogue classes, typically observed in biological samples, while still penalising chemically heterogeneous families; stricter thresholds (≥ 0.9) would over-reject biologically plausible mixed families, whereas more permissive thresholds (≤ 0.5) would inflate apparent coherence.

The component consistency ratio is then defined as:

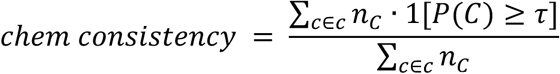

where C is the set f all network components, and 1[⋅] is the indicator function, equal to 1 if the condition inside holds and 0 otherwise. Because each component is weighted by its size *n*_C_, larger consistent families contribute proportionally more to the metric than smaller ones. This metric thus provides a direct measure of the network’s ability to group molecules into chemically coherent families. Metrics were computed on the subset of nodes with available structural annotations; unannotated nodes were excluded from chemistry-based metrics.

#### Rationale for excluding AMI

Adjusted mutual information is a standard measure of agreement between two partitions, and we initially evaluated it alongside the per-component chemistry metrics. However, molecular networks do not partition compounds by chemical class: a single class is routinely split across several disconnected molecular families and adducts, or in-source fragments of the same compound frequently appear as separated nodes. AMI treats these splits as disagreement with the class labels, whereas the consistency metric evaluates each family independently against a purity threshold (≥ 0.7), so a class distributed across multiple pure families still scores well. Including AMI in the composite Z-score degraded the robustness of the top-ranked configurations across datasets, and we therefore report only the per-component chemistry metrics. The results of the comparison of with and without AMI can be found in the arteMIS Zenodo repository (see Data Availability section).

#### Selection of representative configurations

After computing the 100 networks per dataset and calculating the topology and chemistry performance metrics. To identify representative parameter settings for downstream robustness analyses, we applied a multi-criteria ranking approach based on a composite standardized score. Within each dataset × similarity score combination, selected evaluation metrics were standardized across the sampled networks using z-score transformation. The unweighted normalized Z was adopted as the default so that topology and chemistry criteria contribute equally in the absence of a study-specific objective, avoiding researcher bias in the ranking. arteMIS supports user-defined weights so that metrics relevant to a given task (e.g. chemistry metrics for annotation-focused studies, topology for connectivity) can be prioritised; all analyses in this work use the unweighted form.

Metrics for which higher values indicated more desirable network behaviour (high intra-similarity and high chemical consistency) were assigned positive contributions, whereas the high coefficient of node degree variation, number of isolated nodes and Gini coefficient were defined as undesirable and contributed negatively after standardization:

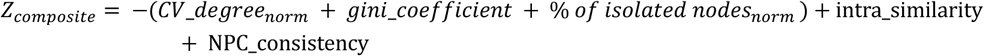

The composite score was calculated as the unweighted sum of these signed standardized metrics, such that higher values correspond to networks that are better connected, less fragmented, and more chemically coherent. Representative top-performing and worst-performing configurations were then defined as those with the highest and lowest composite scores, respectively, while the default configuration corresponds to the current (July 2026) standard values of the GNPS2 for the parameters: cut-off 0.7, maximum links 10 and maximum component size 100.

#### Robustness analysis under subsampling

To evaluate how sensitive the reconstructed molecular networks are under dataset perturbation, we applied a bootstrap-based resampling procedure. For each dataset and score, we used the top-3 configurations obtained in Section 2., for each we applied the subsampling, we computed B=1000 replicates, each retaining a fraction of 85% of the spectra sampled without replacement (meaning a spectrum is a unique sample). For every B replicate, the corresponding submatrix of pairwise similarity scores was extracted and a molecular network was reconstructed. The 85% fraction was chosen to approximate between run variability in feature detection observed across technical replicates, where around 5-20% of features are not consistently detected across acquisitions of the same sample; a higher fraction would underestimate variability, while less node retaining would exceed what is realistically observed between experiments^33,34^.

We computed the same the process for the default configuration (score cut-off = 0.7, maximum of links = 10, maximum component size = 100 and 6 minimum matching peaks). For each configuration, network construction across the B replicates was compared against the reference network using two complementary metrics that capture different aspects of network stability: edge reproducibility and node isolation probability. The results were reported as the average and standard deviation for the top-3 configurations (each represents a different set of parameters) and no average and standard deviation for the unique default configuration.

**Table 7.**
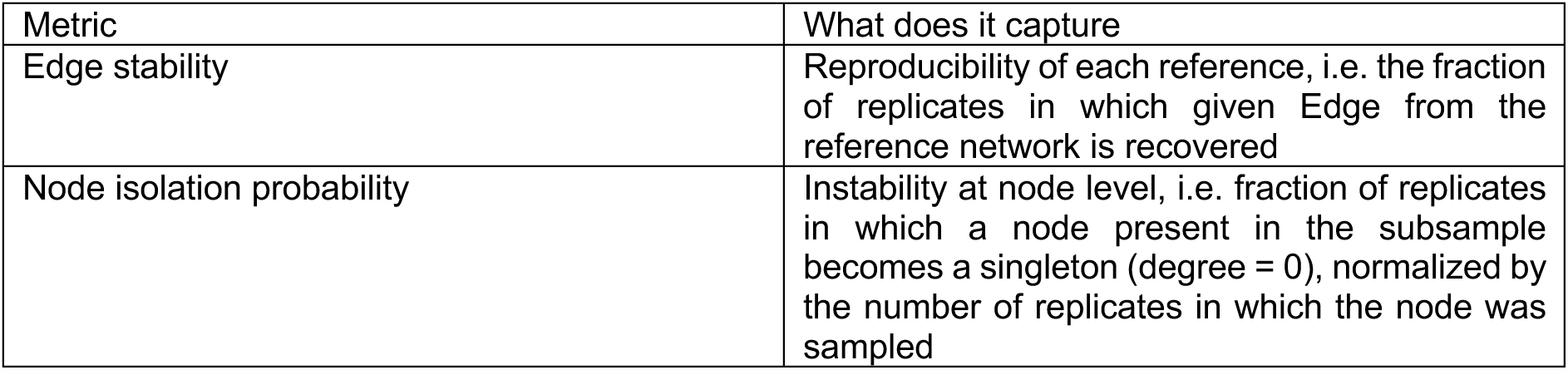
Robustness metric concept and description used.

### Edge stability (Edge reproducibility)

Edge stability quantifies how consistently edges observed in the original network are reproduced across bootstrap replicates.

Let G^(0)^=(V^(0)^,E^(0)^) denote the network constructed using the full dataset, and G^(b)^ = (V^(b)^, E^(b)^), the network obtained from bootstrap replicate *b ε* {1,…*B*}. For each edge *e=(u,v)∈ E^(0)^*, edge stability is defined as the fraction of replicates in which the edge is recovered, conditional on both nodes being retained in the subsample:

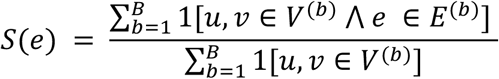

### Node isolation probability

Node isolation probability measures the tendency of nodes to become singletons under resampling.

For each node n and bootstrap replicates G_1_,….,G_M_, let 1[*n* ∈ *V*(*G_i_*)] indicate that n is present in replicate I, and deg_Bi_(n) its degree in that replicate. The node isolation probability is:

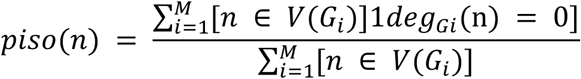

Higher values indicate networks that frequently isolate nodes under dataset perturbation.

#### Case Study seed-mode

A mixture of features from 6 actinomycete strains (*Streptomyces, Microbispora, Actinoplanes, Dactylosporangium, Planomonospora*, and *Micromonospora*) was retrieved from NAICONS collection. The experimental files were preprocessed using mzmine^56^ (version 3.9.0), and the batch file can be found in our Zenodo link. After mzmine, 10,326 features were retained in a mgf file (see Data Availability). From these mgf, Modified Cosine with a threshold of 0.7 was computed against the golden_library from NAICONS, that comprises also authentic standards, the ID to the MGF from NAICONS library is reported in the Supplementary Table 6. along with the spectral scores, features, retention time and more metadata information. These SMILES annotations serve as our ground truth for our seed-annotated compounds. Later, based on these SMILES, using MS2Query we retrieved the NPClassifier pathway_level classification to use later.

The mgf was the input for arteMIS framework, were first we pre-processed the spectra with matchms (version 0.31.0) by applying DEFAULT_FILTERS + CLEAN_PEAKS for a general cleaning, modified cosine score was computed, with a tolerance of 0.02. After the scores are computed, arteMIS generates the Latin Hypercube Sampling. In this case, for cut-off the range was between 0.55 to 0.88, maximum component size from 50 to 200 and maximum links from 5 to 50. After, the 100 networks were computed, for each topology (coefficient of variation of node degree, % isolated nodes and Gini coefficient) and chemistry (intra-similarity and NPClassifier pathway level consistency) were calculated. Afterwards, a z-composite score was generated for the top1, 2 and 3 networks. A default network was generated to be compared against using the GNPS default settings: cut-off = 0.7, maximum component size = 100 and maximum links. = 10. These networks were then evaluated with a pipeline that generates a comparison table between networks and next generates a table to track the annotated features and the molecular families they are part of (if they grow in the top network, shrink or stayed the same), code available, see Data – Availability.

#### Case Study target class-mode

We used the dataset published by Pfütze et al. (2024)^39^. The.mgf file was the input for arteMIS framework, were we first we preprocessed the spectra with matchms (version 0.31.0) by applying DEFAULT_FILTERS as described before, for a general cleaning, leaving 2,225 spectra after cleaning. Modified cosine score were computed with matchms (development branch), using a tolerance of 0.05. arteMIS was employed to generate 100 networks through Latin Hypercube Sampling of the parameter space: maximum component size [30-300], maximum links [5-50], cut-off [0.6-0.8] and matching peaks [2,10]. The calculated discrepancy was 0.005. To complement annotations derived from matches to isolated reference compounds, the same.mgf file was processed with MS2Query to retrieve SMILES predictions for previously unannotated nodes.

For the targeted-class mode, labels were constructed from the default network as follows: all nodes belonging to a connected component that contained at least one annotated melleolide were designated as “melleolide-like”, while all remaining nodes were designated as “others”. These binary labels were then used to guide the optimisation target, replacing the global chemical evaluation metric used in default arteMIS, while minimising their fragmentation across disconnected components. The parameter combination that best satisfied this aim across 100 sampled networks was retained as the optimized configuration.

## Data and Code Availability Statement

The arteMIS codebase is publicly available in our GitHub repository (https://github.com/rtlortega/arteMIS). The documentation is available in the README.md file for installation and execution. All materials needed to reproduce this study are archived on Zenodo (10.5281/zenodo.21622909), including the raw data for the libraries and one of the case studies, links to the Spec2Vec and MS2DeepScore models, the scripts and notebooks used to generate the results and figures, and the results themselves.

## Author contributions

L.R.T.-O., E.C.-G., F.H., and J.J.J.vdH. conceived the concept behind arteMIS. L.R.T.-O. designed and implemented the computational workflow, with E.C.-G. contributing to the conceptualisation and early implementation of the composite-score and target-mode modules, as well as to code testing. F.H. provided the initial code base for network construction. L.R.T.-O. performed the benchmarking analyses across datasets and scoring methods, applied arteMIS to the actinobacteria dataset, and analysed the results; E.C.-G. led the melleolide case study. M.Si. and M.So. generated the experimental data for the actinobacteria case study, performed preprocessing, and delivered the feature table. L.R.T.-O. lead the writing part of the manuscript, with input from E.C.-G. on the melleolide case study; E.C.-G., F.H., M.Si., M.So., and J.J.J.vdH. revised the manuscript. J.J.J.vdH. supervised the project, contributed to the critical interpretation of the results, and participated in manuscript editing. All authors reviewed and approved the final version.

## Acknowledgements

L.R.T.-O. and J.J.J.vdH. gratefully recognize the Marie Skłodowska-Curie grant under the European Union’s Horizon Europe programme MAGiC-MOLFUN (grant no. 101072485).

