## Supplementary Information for "Beyond the Default: Optimizing Molecular Networking with arteMIS"

### Supplementary Figures

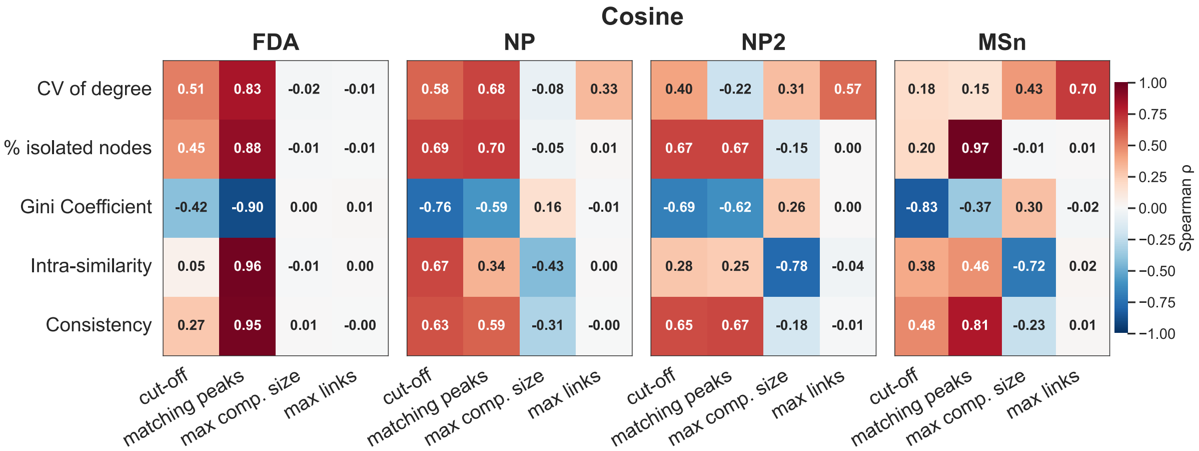

**Supplementary Figure 1.** Pearson Correlation in Cosine-based molecular networks between the parameters: cut-off, matching peaks, maximum component size and maximum links against our different metrics: coefficient of variation of node degree, percentage of isolated nodes, Gini-coefficient, intra-similarity and consistency.

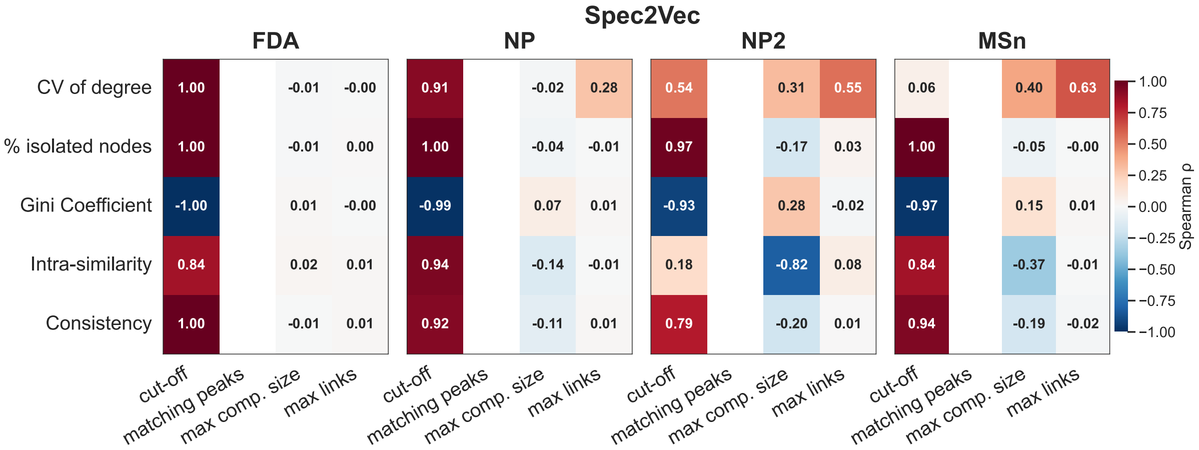

**Supplementary Figure 2.** Pearson Correlation in Spec2Vec[1]- based molecular networks between the parameters: cut-off, matching peaks, maximum component size and maximum links against our different metrics: coefficient of variation of node degree, percentage of isolated nodes, Gini-coefficient, intra-similarity and consistency.

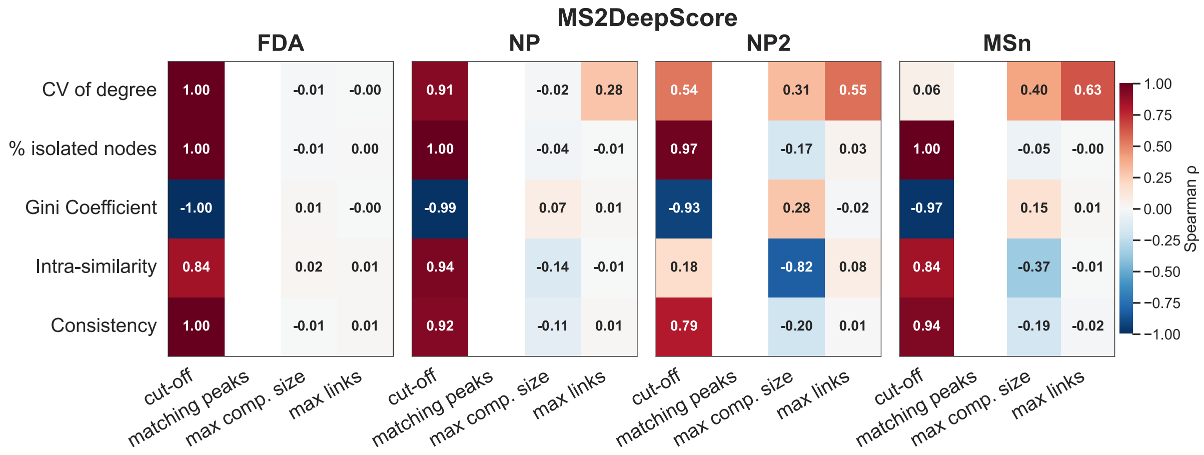

**Supplementary Figure 3.** Pearson Correlation in MS2DeepScore[2]-based molecular networks between the parameters: cut-off, matching peaks, maximum component size and maximum links against our different metrics: coefficient of variation of node degree, percentage of isolated nodes, Gini-coefficient, intra-similarity and consistency.

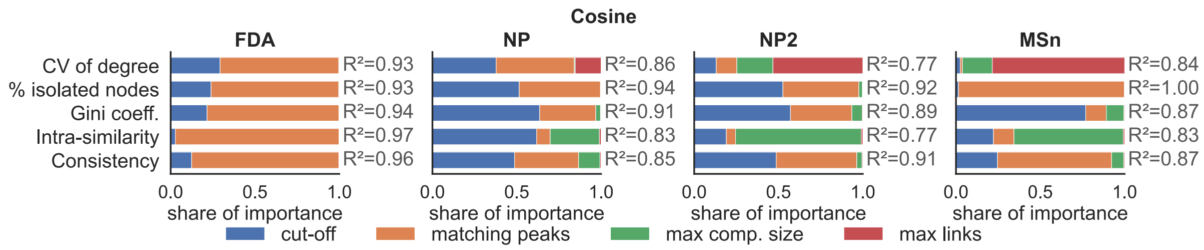

**Supplementary Figure 4.** Random-forest feature (parameter) importances (Cosine-based molecular networks) for the different metrics: coefficient of variation of node degree, percentage of isolated nodes, Gini-coefficient, intra-similarity and consistency.

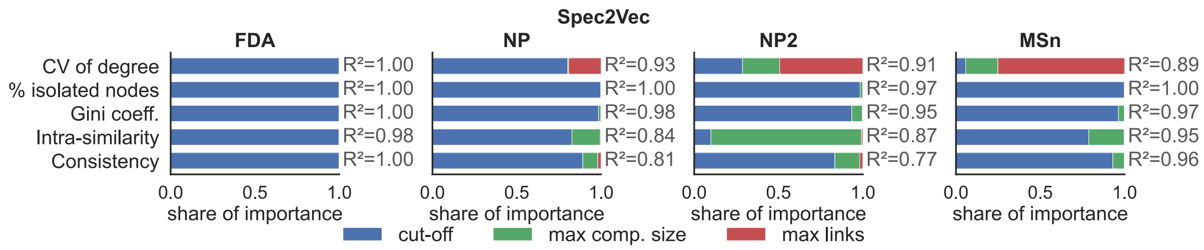

**Supplementary Figure 5.** Random-forest feature (parameter) importances (Spec2Vec- based molecular networks) for the different metrics: coefficient of variation of node degree, percentage of isolated nodes, Gini-coefficient, intra-similarity and consistency.

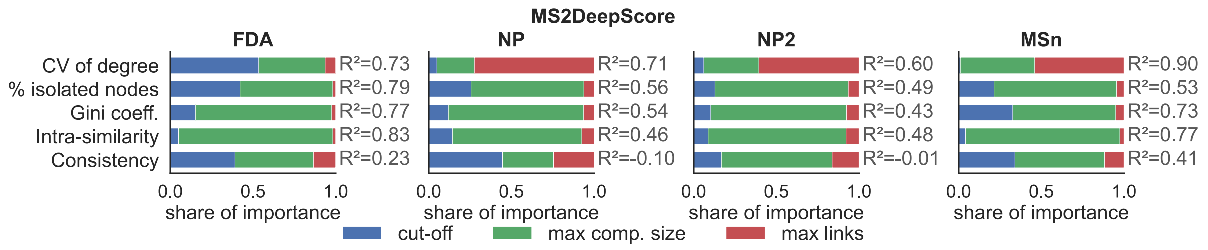

**Supplementary Figure 6.** Random-forest feature (parameter) importances (MS2DeepScore)-based molecular networks for the different metrics: coefficient of variation of node degree, percentage of isolated nodes, Gini-coefficient, intra-similarity and consistency.

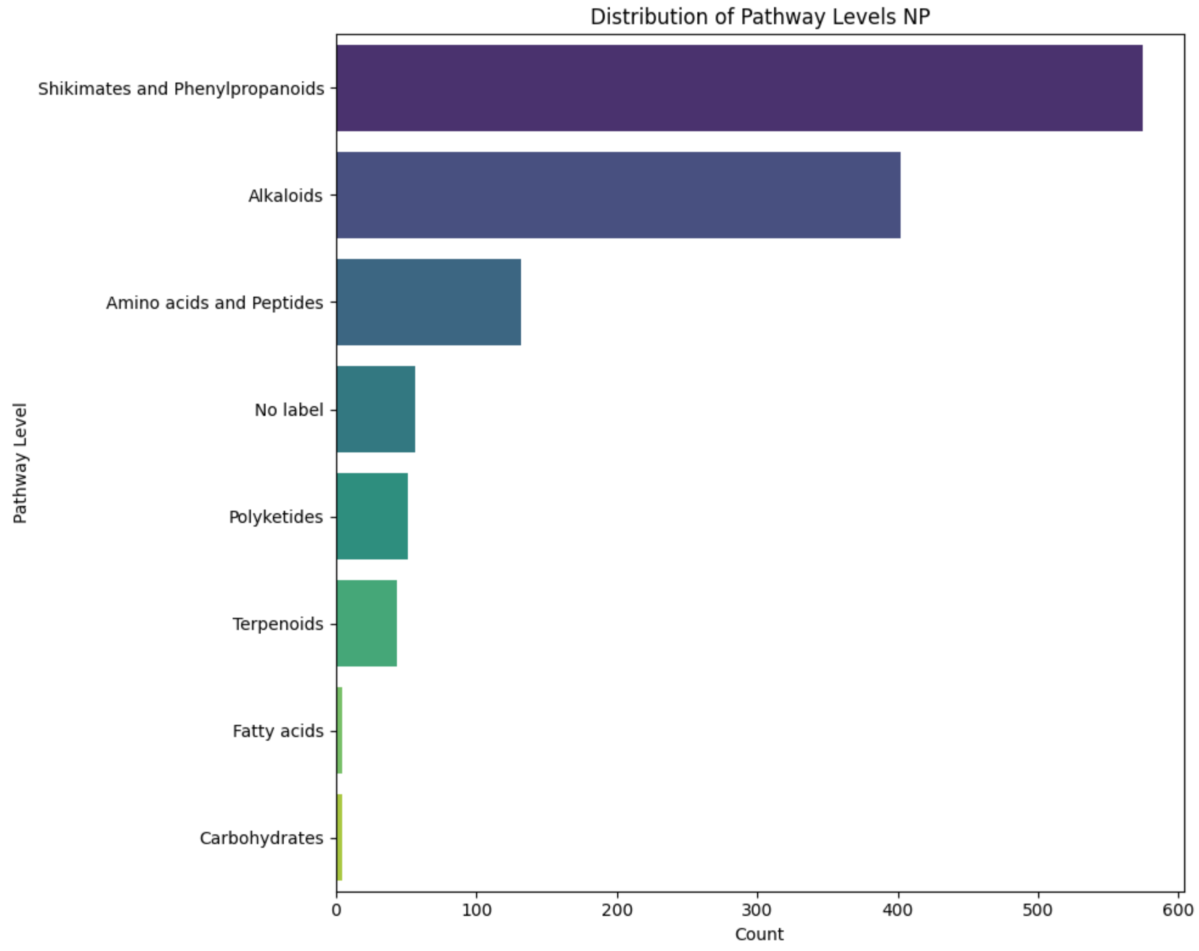

**Supplementary Figure 7.** Chemical pathway level distribution calculated using MS2Query[3] (which retrieves it from NPClassifier[4]) from the NP-dataset (n= 1267).

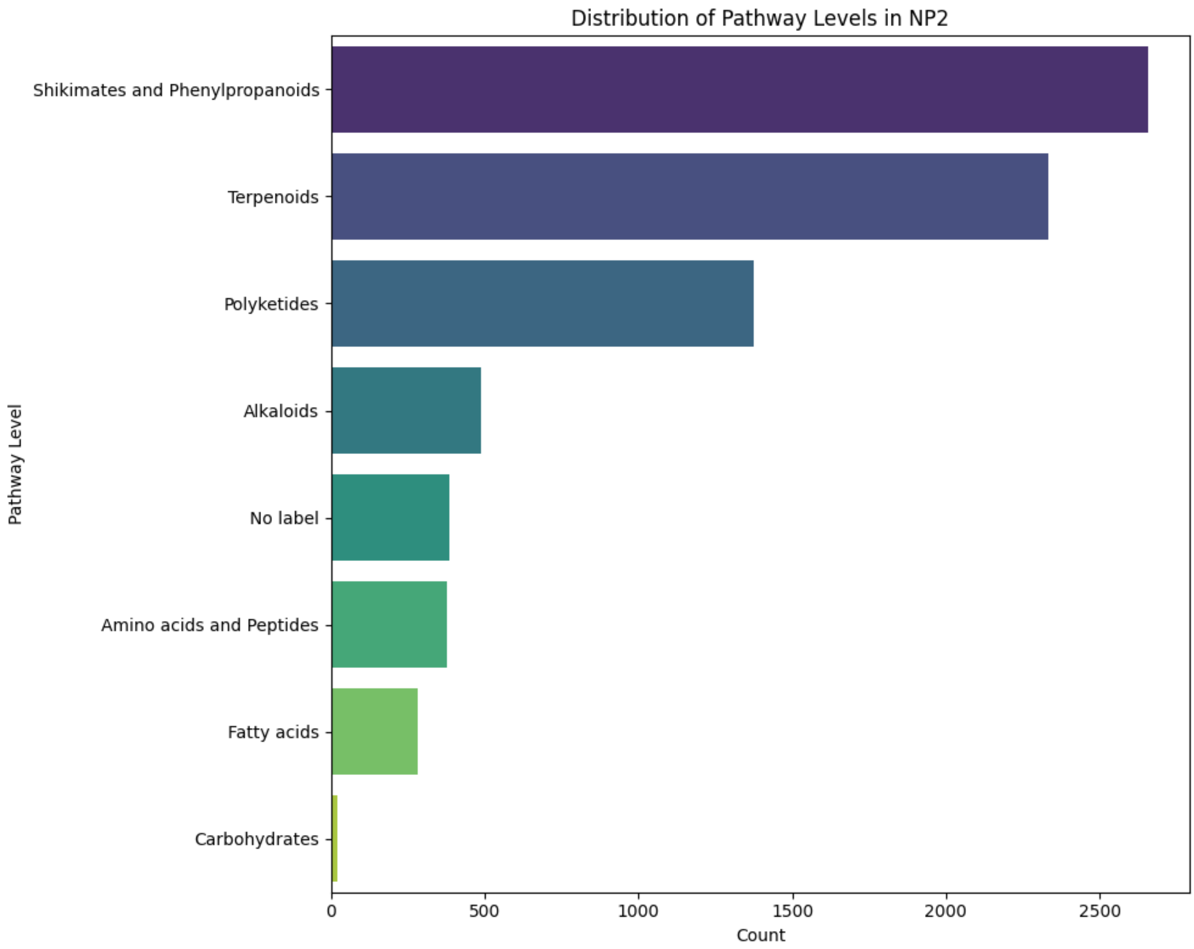

**Supplementary Figure 8.** Chemical pathway level distribution calculated using MS2Query (which retrieves it from NPClassifier[4]) from the NP2-dataset (n=7915).

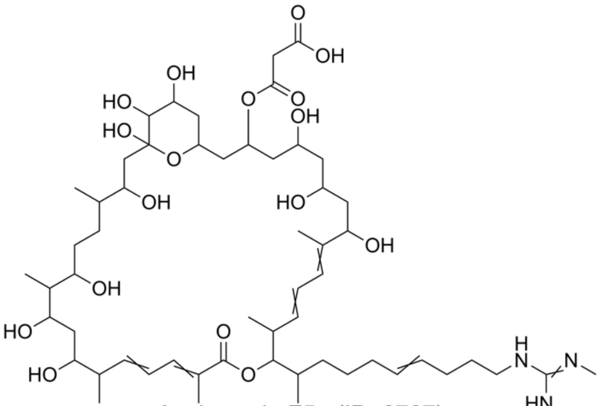

Azalomycin F5a (ID: 6767)

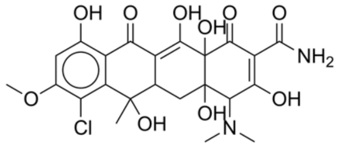

Nigericin like (ID: 8481)

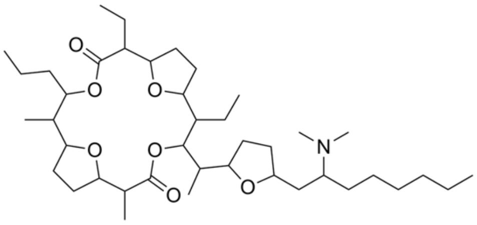

Pamamycin-691 (ID: 8584)

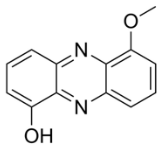

1-1-hydroxy-6-methoxy-phenazine like (ID: 6767)

**Deferroxamines**

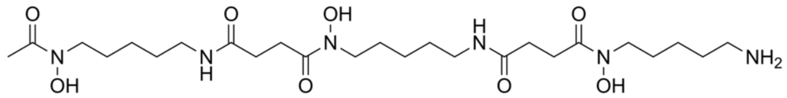

Deferroxamine (ID: 1899)

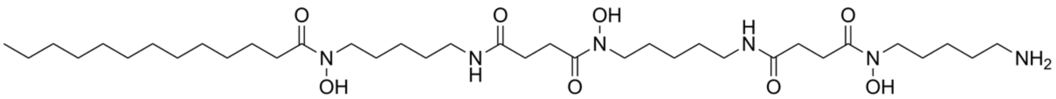

Acyl-Deferroxamine_C12 (ID: 6652 and 6651)

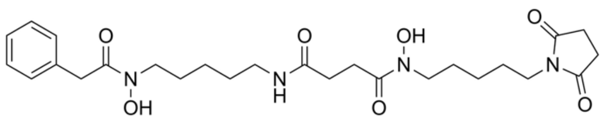

Terragine A (ID: 6652 and 6651)

**Supplementary Figure. 9.** Seed-mode case study singleton rescued. Features that were singletons in default network but were re-joining other families in top1.

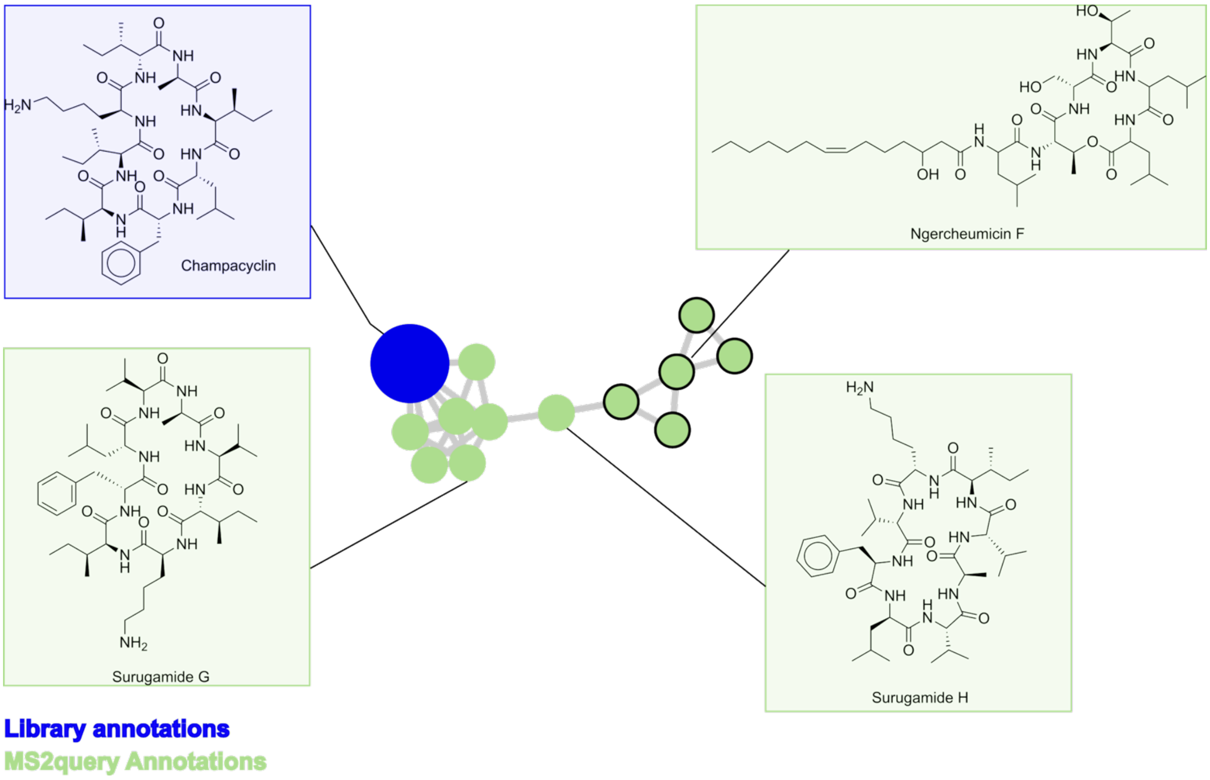

**Supplementary Figure 10.** Seed-mode case study champacyclin example. This is the molecular family in top1. In blue the known compound (champacyclin), in green the analogues found by MS2Query, all “oligopeptides”. Some examples are shown, where they share a cyclised octapeptide scaffold.

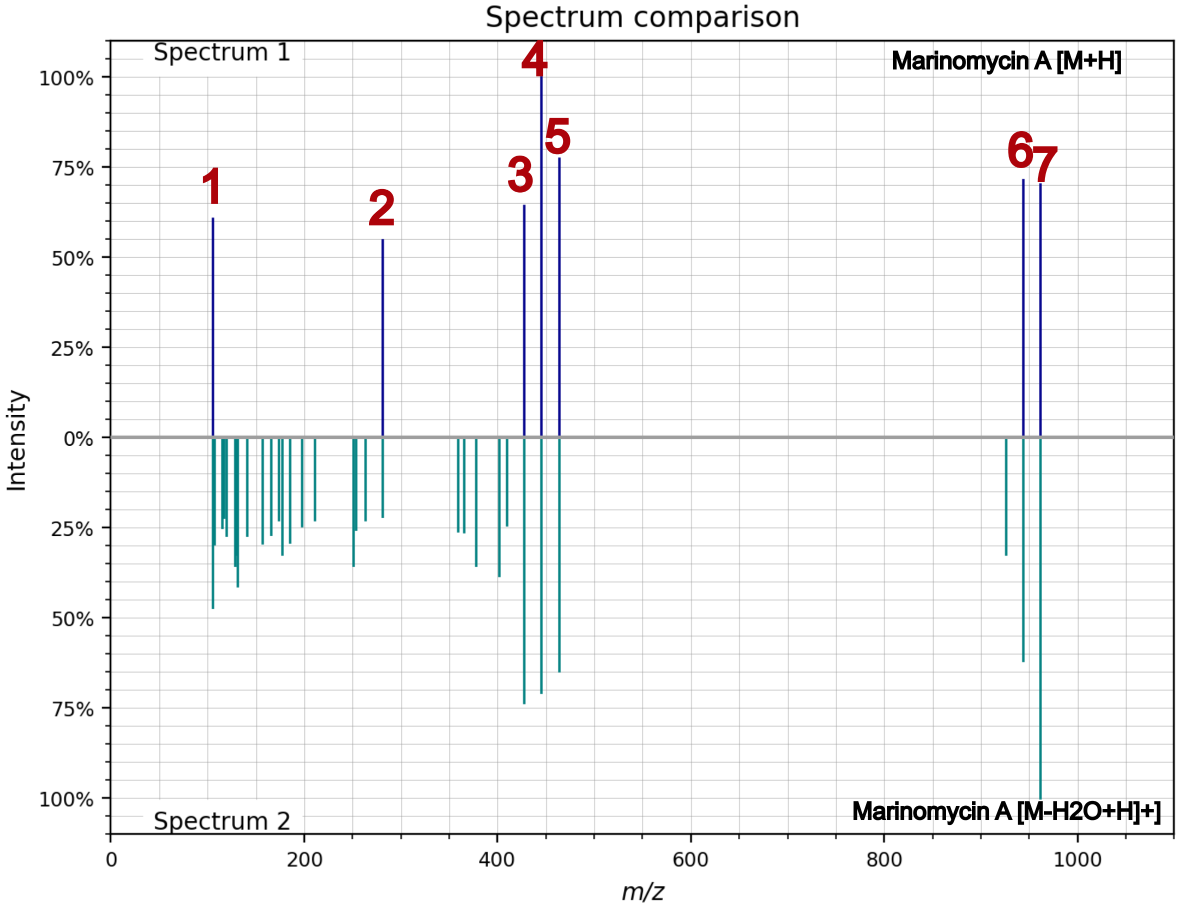

**Supplementary Figure 11.** Seed-mode case study, splitting of compounds that should be together in top1. On top, marinomycin A (node ID: 7366, adduct [M+H]+) against the sample molecule different adduct (node ID: 7365, adduct [M-H2O+H]+). There are 7 matching peaks, the connecting is not retained for top1, since the minimum is 8 peaks.

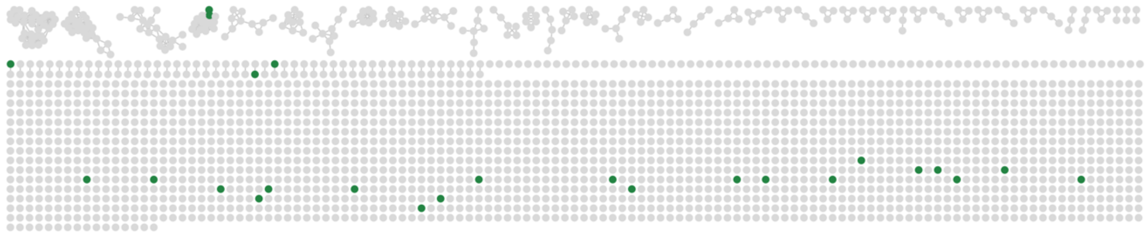

**Supplementary Figure 12.** Global-mode networks for melleolide case study. In green the melleolide standards. Green dots indicate melleolide standards, majorly spotted in different molecular families

#### Supplementary Tables

**Supplementary Table 1.** Results of discrepancy and running times of the benchmarked dataset used. The experiments were conducted on an Apple MacBook Pro (model identifier: Mac16,1), equipped with an Apple M4 chip with 10 CPU cores and 16 GB of unified memory.

| total spectra | no. spectra after cleaning | cut off | max comp size | max links | matching peaks | Discrepancy Cosine-based | Discrepancy ML-based | Time (s) Cosine-based | Time (s) ML-based |
|---|---|---|---|---|---|---|---|---|---|
| 656 | 606 | [0.55-0.8] | [10, 100] | [5,20] | [2,8] | 0.00053 | 0.0002 | 14,5 | 27,3 |
| 1267 | 1226 | [0.6-0.8] | [30, 300] | [5,50] | [2,10] | 0.00053 | 0.0002 | 53,3 | 83,3 |
| 7915 | 7250 | [0.6-0.85] | [50-400] | [8,100] | [2,10] | 0.00053 | 0.000 | 1518,0 | 2354,9 |
| 15920 | 12929 | [0.65-0.85] | [50-500] | [8,100] | [3,10] | 0.00053 | 0.0002 | 5093,7 | 8305,4 |

**Supplementary Table 2.** Z-score statistical results of the 100-networks computed in Section 2. per score and dataset.

| Score | Dataset | Minimum | Median | Maximum |
|---|---|---|---|---|
| Cosine | FDA | -2.37 | 0.34 | 1.88 |
| MS2DeepScore | FDA | -4.03 | 0.05 | 5.08 |
| Modified Cosine | FDA | -4.06 | 0.25 | 4.01 |
| Spec2Vec | FDA | -2.22 | 0.19 | 1.96 |
| Cosine | NP | -3.07 | -0.17 | 3.44 |
| MS2DeepScore | NP | -4.58 | 0.35 | 3.8 |
| Modified Cosine | NP | -4.79 | 0.09 | 4.6 |
| Spec2Vec | NP | -4.1 | 0.5 | 2.83 |
| Cosine | NP2 | -4.44 | 0.08 | 5.17 |
| MS2DeepScore | NP2 | -3.82 | -0.24 | 4.77 |
| Modified Cosine | NP2 | -5.19 | 0.15 | 3.17 |
| Spec2Vec | NP2 | -3.46 | -0.13 | 5.6 |
| Cosine | MSn | -5.28 | -0.24 | 6.53 |
| MS2DeepScore | MSn | -5.17 | -0.09 | 6.03 |
| Modified Cosine | MSn | -6.68 | -0.16 | 8.05 |
| Spec2Vec | MSn | -4.89 | 0.43 | 5.28 |

**Supplementary Table 3.** Robustness of top-ranked and default configurations across datasets and spectral similarity scores. Top-configuration parameter values and metrics are reported as mean ± SD across the three top-ranked configurations (n = 3); default is the GNPS configuration (single values). Per-configuration metrics were computed over B = 1000 bootstrap replicates (85% subsampling). P_iso = mean node isolation probability; Edge instability = fraction of reference edges recovered in fewer than 95% of bootstrap replicates. Min matching peaks is blank for MS2DeepScore and Spec2Vec as the parameter is not applicable to these scores.

| Dataset | Score | Config | Max comp size | Cut-off | Max links | Min matching peaks | P_iso | Edge instability (<0.95) |
|---|---|---|---|---|---|---|---|---|
| FDA-dataset | Cosine | default | 100 | 0.7 | 10 | 6 | 0.889 | 0 |
| FDA-dataset | Cosine | top (n=3) | 68 ± 20 | 0.58 ± 0.01 | 13 ± 4 | 7.3 ± 0.6 | 0.880 ± 0.013 | 0 |
| FDA-dataset | Modified Cosine | default | 100 | 0.7 | 10 | 6 | 0.71 | 0 |
| FDA-dataset | Modified Cosine | top (n=3) | 16 ± 5 | 0.73 ± 0.06 | 10 ± 4 | 4.7 ± 1.2 | 0.680 ± 0.020 | 0 |
| FDA-dataset | MS2DeepScore | default | 100 | 0.7 | 10 | - | 0.482 | 0 |
| FDA-dataset | MS2DeepScore | top (n=3) | 13 ± 2 | 0.66 ± 0.05 | 16 ± 2 | - | 0.567 ± 0.012 | 0.032 ± 0.020 |
| FDA-dataset | Spec2Vec | default | 100 | 0.7 | 10 | - | 0.841 | 0 |
| FDA-dataset | Spec2Vec | top (n=3) | 62 ± 19 | 0.72 | 10 ± 3 | - | 0.858 | 0 |
| NP-dataset | Cosine | default | 100 | 0.7 | 10 | 6 | 0.372 | 0 |
| NP-dataset | Cosine | top (n=3) | 55 ± 18 | 0.73 ± 0.02 | 12 ± 10 | 6.7 ± 2.5 | 0.410 ± 0.052 | 0 |
| NP-dataset | Modified Cosine | default | 100 | 0.7 | 10 | 6 | 0.274 | 0.003 |
| NP-dataset | Modified Cosine | top (n=3) | 146 ± 60 | 0.69 ± 0.05 | 10 ± 1 | 4.3 ± 1.2 | 0.259 ± 0.003 | 0.010 ± 0.006 |
| NP-dataset | MS2DeepScore | default | 100 | 0.7 | 10 |  | 0.205 | 0.01 |
| NP-dataset | MS2DeepScore | top (n=3) | 188 ± 121 | 0.69 ± 0.08 | 6 |  | 0.197 ± 0.020 | 0.010 ± 0.008 |
| NP-dataset | Spec2Vec | default | 100 | 0.7 | 10 |  | 0.464 | 0 |
| NP-dataset | Spec2Vec | top (n=3) | 81 ± 52 | 0.71 ± 0.04 | 6 ¬± 2 |  | 0.478 ± 0.054 | 0.003 ± 0.005 |
| NP2-dataset | Cosine | default | 100 | 0.7 | 10 | 6 | 0.348 | 0.001 |
| NP2-dataset | Cosine | top (n=3) | 55 ± 5 | 0.78 ± 0.07 | 47 ± 15 | 6.0 ± 1.7 | 0.431 ± 0.077 | 0.002 ± 0.001 |
| NP2-dataset | Modified Cosine | default | 100 | 0.7 | 10 | 6 | 0.385 | 0.008 |
| NP2-dataset | Modified Cosine | top (n=3) | 87 ± 16 | 0.71 ± 0.06 | 20 ± 17 | 8.3 ± 1.2 | 0.384 ± 0.016 | 0.010 ± 0.007 |
| NP2-dataset | MS2DeepScore | default | 100 | 0.7 | 10 | - | 0.416 | 0.001 |
| NP2-dataset | MS2DeepScore | top (n=3) | 67 ± 10 | 0.68 ± 0.06 | 55 ± 40 | - | 0.417 ± 0.004 | 0.001 ± 0.001 |
| NP2-dataset | Spec2Vec | default | 100 | 0.7 | 10 | - | 0.424 | 0.01 |
| NP2-dataset | Spec2Vec | top (n=3) | 59 ± 9 | 0.75 ± 0.10 | 60 ± 27 | - | 0.508 ± 0.130 | 0.002 ± 0.000 |
| MSn-dataset | Cosine | default | 100 | 0.7 | 10 | 6 | 0.083 | 0.019 |
| MSn-dataset | Cosine | top (n=3) | 57 ± 6 | 0.79 ± 0.06 | 47 ± 15 | 6.0 ± 1.7 | 0.104 ± 0.060 | 0.003 ± 0.003 |
| MSn-dataset | Modified Cosine | default | 100 | 0.7 | 10 | 6 | 0.081 | 0.035 |
| MSn-dataset | Modified Cosine | top (n=3) | 57 ± 6 | 0.79 ± 0.06 | 47 ± 15 | 6.0 ± 1.7 | 0.099 ± 0.058 | 0.003 ± 0.002 |
| MSn-dataset | MS2DeepScore | default | 100 | 0.7 | 10 | - | 0.04 | 0.022 |

| | | | | | | | | |
|---|---|---|---|---|---|---|---|---|
| MSn-dataset | MS2DeepScore | top (n=3) | 97 ± 57 | 0.84 ± 0.01 | 61 ± 35 | - | 0.053 ± 0.005 | 0.001 ± 0.001 |
| MSn-dataset | Spec2Vec | default | 100 | 0.7 | 10 | - | 0.047 | 0.006 |
| MSn-dataset | Spec2Vec | top (n=3) | 246 ± 182 | 0.83 ± 0.02 | 17 ± 10 | - | 0.089 ± 0.009 | 0.000 ± 0.001 |

**Supplementary Table 4.** Seed-mode case study results. Changes in molecular family composition default vs. top.

| Feature_id | Compound_name | Default_size | Top1_size |
|---|---|---|---|
| 789 | bacilysin | 1 | 1 |
| 1719 | Pyridindolol | 1 | 1 |
| 2366 | FUTALOSINE | 1 | 1 |
| 2596 | Pantethine | 1 | 1 |
| 2712 | dehydroxynocardamine | 1 | 1 |
| 2736 | dehydroxynocardamine | 1 | 1 |
| 2947 | nocardamin | 1 | 1 |
| 4062 | N-Acetyl-Acyl-Desferrioxamine_C8_Keto | 1 | 1 |
| 4416 | colibrimycin B1 | 1 | 1 |
| 4460 | Chymostatinol_A | 1 | 1 |
| 4583 | Daidzein | 1 | 1 |
| 4768 | NAI-107_F1 | 1 | 1 |
| 4832 | NAI-107_F2 | 1 | 1 |
| 4904 | Lydicamycin | 1 | 1 |
| 5133 | Indolokine_A4 | 1 | 1 |
| 5176 | NAI-107_A1 | 1 | 1 |
| 5217 | NAI-107_A2 | 1 | 1 |
| 5279 | colibrimycin A1 demethyl-analogue | 1 | 1 |
| 5364 | NAI-107_A0 | 1 | 1 |
| 5538 | colibrimycin A1 | 1 | 1 |
| 5569 | Furaquinocin_I | 1 | 1 |
| 5610 | TPU-0037-C | 1 | 1 |
| 5779 | Propeptin | 1 | 1 |
| 5907 | Dactylosporolide_A | 1 | 1 |
| 6020 | 5,6-Dihydroxyphenazin-1(5H)-one | 1 | 1 |
| 6047 | PHENAZINE | 1 | 1 |
| 6226 | N-Acetyl-Acyl-Desferrioxamine_C9 | 1 | 1 |
| 6371 | COPROPORPHYRIN_III | 1 | 1 |
| 6378 | COPROPORPHYRIN_III | 1 | 1 |
| 6416 | 1,6-Dihydroxyphenazine | 1 | 1 |
| 6493 | AZALOMYCIN_F-5A | 1 | 1 |
| 6527 | colibrimycin C6 | 1 | 1 |
| 6559 | Surugamide_A | 1 | 1 |

| | | | |
|---|---|---|---|
| 6566 | Champacyclin | 1 | 1 |
| 6767 | Azalomycin F5a | 1 | 2 |
| 6895 | Alteramide A | 1 | 1 |
| 6906 | Alteramide A | 1 | 1 |
| 7160 | Ikarugamycin_epoxide | 1 | 1 |
| 7334 | Maltophilin | 1 | 1 |
| 7411 | MARINOMYCIN_A | 1 | 1 |
| 7783 | PIERICIDIN_A1 | 1 | 1 |
| 7840 | Antimycin A19 | 1 | 1 |
| 7903 | 21-hydroxy oligomycin C | 1 | 1 |
| 7904 | 21-hydroxy oligomycin C | 1 | 1 |
| 8481 | Nigericin | 1 | 1 |
| 8491 | TETRANACTIN | 1 | 1 |
| 1962 | desferrioxamine G | 1 | 1 |
| 8721 | Nigericin like | 1 | 2 |
| 6767 | "1-Hydroxy-6-methoxy-phenazine;_6-Methoxy-1-phenazinol" | 1 | 4 |
| 1899 | Deferoxamine | 1 | 4 |
| 6651 | Acyl-Desferrioxamine_C12 | 1 | 66 |
| 6652 | Acyl-Desferrioxamine_C12 | 1 | 75 |
| 8584 | Pamamycin-691 | 1 | 109 |
| 1889 | Deferoxamine | 1 | 166 |
| 4217 | Terragine_A | 1 | 166 |
| 644 | ALTEMICIDIN | 2 | 1 |
| 2729 | dehydroxynocardamine | 2 | 2 |
| 2949 | nocardamin | 2 | 2 |
| 4132 | N-Acetyl-Acyl-Desferrioxamine_C8_Keto | 2 | 2 |
| 4589 | Chymostatinol_A | 2 | 2 |
| 4724 | TPU-0037-A_Lydicamycin | 2 | 2 |
| 6234 | N-Acetyl-Acyl-Desferrioxamine_C9 | 2 | 2 |
| 6552 | Surugamide_A | 2 | 2 |
| 6901 | Alteramide A | 2 | 2 |
| 7158 | Alteramide B | 2 | 2 |
| 8703 | Ornithine-containing lipid_653 | 2 | 2 |
| 6040 | Azalomycin F4a | 2 | 3 |
| 6410 | Azalomycin F4a | 2 | 3 |
| 5519 | colibrimycin A1 | 2 | 53 |
| 5581 | colibrimycin A1 | 2 | 53 |
| 6918 | Alteramide A | 3 | 1 |
| 955 | bacilysin | 3 | 2 |
| 2942 | nocardamin | 3 | 2 |
| 8359 | S-3466 A | 3 | 3 |
| 8444 | S-3466 B | 3 | 3 |

| | | | |
|---|---|---|---|
| 8500 | TETRANACTIN | 3 | 3 |
| 6558 | Surugamide_A | 3 | 4 |
| 4602 | Abyssomicin D | 3 | 6 |
| 4710 | colibrimycin A1 demethyl-analogue | 3 | 53 |
| 6525 | colibrimycin C6 | 3 | 53 |
| 5962 | TPU-0037-B | 4 | 1 |
| 5456 | TPU-0037-A_Lydicamycin | 4 | 3 |
| 5637 | TPU-0037-B | 4 | 3 |
| 6193 | SOYASAPONINE_I | 4 | 4 |
| 4693 | TPU-0037-A_Lydicamycin | 4 | 4 |
| 5612 | TPU-0037-C | 4 | 4 |
| 5638 | TPU-0037-B | 4 | 4 |
| 6046 | Surugamide_F | 4 | 4 |
| 8829 | WA8242A1 | 4 | 4 |
| 892 | Sesbanimide A | 4 | 6 |
| 5625 | NAI-107_B2 | 5 | 5 |
| 6052 | Surugamide_F | 6 | 3 |
| 22 | Exochelin_MS_analogue | 6 | 4 |
| 29 | Exochelin_MS;<NL>N-(dN-Formyl,dN-hydroxy-R-ornithinyl)-beta-alaninyl-dN-hydroxy-R-ornit<NL>-R-allo-threoninyl-dN-hydroxy-S-ornithine | 6 | 4 |
| 6183 | N-Acetyl-Acyl-Desferrioxamine_C9 | 6 | 5 |
| 6316 | Azalomycin F3a | 6 | 9 |
| 1981 | Dibenarthin | 6 | 10 |
| 4298 | Chymostatinol_A | 7 | 7 |
| 4678 | CHYMOSTATINOL_C | 7 | 7 |
| 4895 | TPU-0037-B | 7 | 7 |
| 6557 | Champacyclin | 7 | 13 |
| 4371 | 7-O-methyl-genistein | 8 | 8 |
| 4733 | 7-O-methyl-genistein | 8 | 8 |
| 1446 | amiclenomycin | 8 | 58 |
| 8546 | Ornithine-containing-lipid-like | 9 | 1 |
| 8702 | Ornithine-containing lipid_653 | 9 | 1 |
| 8916 | Ornithine-containing-lipid-like | 9 | 6 |
| 9211 | WA8242A1 | 9 | 6 |
| 2142 | desferrioxamine G | 9 | 8 |
| 5563 | Heptapropylene glycol | 11 | 11 |
| 5272 | CHYMOSTATINOL_C | 12 | 12 |
| 4580 | Daidzein | 13 | 13 |
| 5511 | Genistein | 13 | 13 |
| 7399 | Antimycin A5 | 14 | 14 |
| 7528 | Antimycin A20 | 14 | 14 |
| 7738 | Antimycin A17 | 14 | 14 |

| | | | |
|---|---|---|---|
| 7945 | Antimycin A15 | 14 | 14 |
| 7366 | MARINOMYCIN_A | 18 | 1 |
| 7365 | MARINOMYCIN_A | 18 | 17 |
| 7908 | Nigericin | 19 | 27 |
| 7909 | Nigericin | 19 | 27 |
| 8514 | Nigericin | 19 | 27 |
| 8516 | Nigericin | 19 | 27 |
| 8547 | Ornithine-containing-lipid-like | 24 | 22 |
| 4073 | 9-(4'-aminophenyl)-3,7-dihydroxy-2,4,6-trimethyl-9-oxo-nonoic_acid | 25 | 24 |
| 6133 | Levorin A3 | 25 | 24 |
| 6210 | Levorin A2 | 25 | 24 |
| 2440 | AMASTATIN_B2 | 25 | 25 |
| 1905 | Deferoxamine | 27 | 26 |
| 6631 | Acyl-Desferrioxamine_C12 | 27 | 26 |
| 6177 | SOYASAPONINE_I | 35 | 33 |
| 6482 | Siomycin A | 36 | 39 |
| 6231 | N-Acetyl-Acyl-Desferrioxamine_C9 | 37 | 37 |
| 8337 | Ornithine-containing lipid_597 | 39 | 39 |
| 8412 | Ornithine-containing lipid-like_611 | 39 | 39 |
| 8452 | Ornithine-containing-lipid-like | 39 | 39 |
| 8535 | Ornithine-containing lipid_597 | 39 | 39 |
| 8538 | Ornithine-containing lipid | 39 | 39 |
| 8555 | Ornithine-containing lipid-like_611 | 39 | 39 |
| 8637 | Ornithine-containing-lipid-like | 39 | 39 |
| 8762 | Ornithine-containing lipid | 39 | 39 |
| 9061 | Ornithine-containing lipid_653 | 39 | 39 |
| 9367 | Ornithine-containing lipid | 39 | 39 |
| 5304 | colibrimycin A1 demethyl-analogue | 46 | 53 |
| 5305 | colibrimycin A1 demethyl-analogue | 46 | 53 |
| 5330 | colibrimycin A1 demethyl-analogue | 46 | 53 |
| 5334 | colibrimycin A1 demethyl-analogue | 46 | 53 |
| 5576 | colibrimycin A1 | 46 | 53 |
| 5578 | colibrimycin A1 | 46 | 53 |
| 5598 | colibrimycin A1 | 46 | 53 |
| 2102 | Dibenarthin | 47 | 59 |
| 4333 | GE20372 B | 47 | 59 |
| 4463 | Chymostatinol_A | 47 | 59 |
| 4599 | GE20372 B | 47 | 59 |
| 4923 | Chymostatinol_A | 47 | 59 |
| 5284 | CHYMOSTATINOL_C | 47 | 59 |
| 5354 | Chymostatin_analog_4 | 47 | 59 |
| 6455 | Azalomycin F4a | 47 | 106 |

| | | | |
|---|---|---|---|
| 4267 | Terragine_A | 54 | 2 |
| 2527 | Daidzin | 58 | 32 |
| 2566 | Daidzin | 58 | 32 |
| 3003 | Daidzin | 58 | 32 |
| 3301 | Genistin | 58 | 32 |
| 5588 | indolokine_A5 | 76 | 54 |
| 7775 | PIERICIDIN_A1 | 82 | 4 |
| 888 | Sesbanimide A | 82 | 11 |

**Supplementary Table 5.** Global-mode top configurations for the melleolide case stuy.

| Max Comp Size | Max links | Cut off | Matching peaks | Degree_CV | % isolate nodes | Gini-coef. | Intra-similarity | Consistency | Z-score |
|---|---|---|---|---|---|---|---|---|---|
| 111 | 7 | 0.8 | 3 | 0.767 | 0.840 | 0.116 | 0.120 | 0.974 | 3.206 |
| 144 | 11 | 0.73 | 3 | 0.743 | 0.762 | 0.183 | 0.119 | 0.917 | 2.721 |
| 150 | 6 | 0.66 | 10 | 0.688 | 0.731 | 0.236 | 0.116 | 0.914 | 2.662 |

**Supplementary Table 6.** Annotated compounds for the seed-mode case study

| feature_id | compound_name | adduct | score | precursor_mz | rt | mol_formula | inchi |
|---|---|---|---|---|---|---|---|
| 22 | Exochelin_MS_analogue | [M+2H]+2 | 0.913 | 333.6745911 | 0.6938997 | C25H47N9O12 | None |
| 29 | Exochelin_MS;<NL>N-(dN-Formyl,dN-hydroxy-R-ornithinyl)-beta-alaninyl-dN-hydroxy-R-ornit<NL>-R-allo-threoninyl-dN-hydroxy-S-ornithine | [M+2H]+2 | 0.953 | 305.1635437 | 0.55372924 | C23H44N8O11 | AHHBHDOBNNJFAK |
| 644 | ALTEMICIDIN | [M+H]+ | 0.934 | 377.1123352 | 1.0889416 | C13H20N4O7S | VZRFZUPFQKSXPV-VPFIQFBESA-N |
| 789 | bacilysin | [M-H2O+H]+ | 0.905 | 253.1183624 | 1.3640732 | C12H18N2O5 | XFOUAXMJRHNTOP |
| 888 | Sesbanimide A | [M+H]+ | 0.992 | 328.1389008 | 1.5251907 | C15H21NO7 | ULQATHQJWVNXEJ-UHFFFAOYAF |
| 892 | Sesbanimide A | [M-H2O+H]+ | 0.98 | 310.1283569 | 1.3669575 | C15H21NO7 | ULQATHQJWVNXEJ-UHFFFAOYAF |
| 955 | bacilysin | [M-H2O+H]+ | 0.885 | 253.1183624 | 1.3640732 | C12H18N2O5 | XFOUAXMJRHNTOP |
| 1446 | amiclenomycin | [M+H]+ | 0.949 | 197.1283875 | 3.6329668 | C10H16N2O2 | LAJWZJCOWPUSOA |
| 1719 | Pyridindolol | [M+H]+ | 0.985 | 259.1077118 | 4.13734 | C14H14N2O3 | VAKXHGQDPLEUTH-LBPRGKRZBA |
| 1889 | Deferoxamine | [M+2H]+2 | 0.975 | 281.1838074 | 4.043571 | C25H48N6O8 | UBQYURCVBFRUQT-UHFFFAOYAW |
| 1899 | Deferoxamine | [M+H]+ | 0.997 | 561.3601379 | 4.112325 | C25H48N6O8 | UBQYURCVBFRUQT-UHFFFAOYAW |
| 1905 | Deferoxamine | [M+Na]+ | 0.993 | 583.3419189 | 4.125219 | C25H48N6O8 | UBQYURCVBFRUQT-UHFFFAOYAW |
| 1962 | desferrioxamine G | [M+H]+ | 0.926 | 619.3656921 | 4.249046 | C27H50N6O10 | MIVGUYBAQIHKPJ-UHFFFAOYAW |
| 1981 | Dibenarthin | [M+2H]+2 | 0.968 | 403.1767883 | 4.2929254 | C34H48N10O13 | WUGMTNYESVFCDC-MEVGPWJASA-N |
| 2102 | Dibenarthin | [M+2H]+2 | 0.949 | 403.1767883 | 4.2929254 | C34H48N10O13 | WUGMTNYESVFCDC-MEVGPWJASA-N |
| 2142 | desferrioxamine G | [M+Na]+ | 0.959 | 641.3477173 | 4.6942925 | C27H50N6O10 | MIVGUYBAQIHKPJ-UHFFFAOYAW |
| 2366 | FUTALOSINE | [M+H]+ | 0.943 | 415.1248779 | 4.5116987 | C19H18N4O7 | VEDWXCWBMDQNCV |

| | | | | | | | |
|---|---|---|---|---|---|---|---|
| 2440 | AMASTATIN_B2 | [M+H]+ | 0.955 | 489.2921753 | 4.755074 | C22H40N4O8 | JFROKIOSWBRFPQ |
| 2527 | Daidzin | [M+H]+ | 0.969 | 417.1180522 | 4.63685 | C21H20O9 | KYQZWONCHDNPDP-QNDFHXLGBS |
| 2566 | Daidzin | [M+Na]+ | 0.9 | 439.0995789 | 4.8545184 | C21H20O9 | KYQZWONCHDNPDP-QNDFHXLGBS |
| 2596 | Pantethine | [M+Na]+ | 0.987 | 577.2337036 | 4.8171434 | C22H42N4O8S2 | DJWYOLJPSHDSAL |
| 2712 | dehydroxynocardamine | [M+Na]+ | 0.978 | 607.342041 | 4.805539 | C27H48N6O8 | ABHHIGWFFMCQOC-UHFFFAOYAM |
| 2729 | dehydroxynocardamine | [M+2H]+2 | 0.915 | 293.1836548 | 4.8364153 | C27H48N6O8 | ABHHIGWFFMCQOC-UHFFFAOYAM |
| 2736 | dehydroxynocardamine | [M+H]+ | 0.988 | 585.3606873 | 4.9372234 | C27H48N6O8 | ABHHIGWFFMCQOC-UHFFFAOYAM |
| 2942 | nocardamin | [M+H]+ | 0.995 | 601.3549805 | 5.0566454 | C27H48N6O9 | NHKCCADZVLTPPO-UHFFFAOYAU |
| 2947 | nocardamin | [M+K]+ | 0.983 | 639.31073 | 5.2047915 | C27H48N6O9 | NHKCCADZVLTPPO-UHFFFAOYAU |
| 2949 | nocardamin | [M+Na]+ | 0.997 | 623.3371582 | 5.034436 | C27H48N6O9 | NHKCCADZVLTPPO-UHFFFAOYAU |
| 3003 | Daidzin | [M+Na]+ | 0.905 | 439.0995789 | 4.8545184 | C21H20O9 | KYQZWONCHDNPDP-QNDFHXLGBS |
| 3301 | Genistin | [M+H]+ | 0.97 | 433.1129303 | 5.345604 | C21H20O10 | ZCOLJUOHXJRHDI-IJFYFLOXBC |
| 4062 | N-Acetyl-Acyl-Desferrioxamine_C8_Keto | [M+2H]+2 | 0.818 | 358.2335815 | 6.2815804 | C34H62N6O10 | JROJQIHMGGIKDI |
| 4073 | 9-(4'-aminophenyl)-3,7-dihydroxy-2,4,6-trimethyl-9-oxo-nonoic_acid | [M+H]+ | 0.981 | 338.1960144 | 6.0785003 | C18H27NO5 | UHZAKFDEIOBYGZ |
| 4132 | N-Acetyl-Acyl-Desferrioxamine_C8_Keto | [M+Na]+ | 0.993 | 737.4415283 | 6.41352 | C34H62N6O10 | JROJQIHMGGIKDI |
| 4217 | Terragine_A | [M+H]+ | 0.938 | 519.2813721 | 6.252744 | C26H38N4O7 | OIEMCIKPOVGWLI |
| 4267 | Terragine_A | [M+NH4]+ | 0.946 | 536.3079224 | 6.321573 | C26H38N4O7 | OIEMCIKPOVGWLI |
| 4298 | Chymostatinol_A | [M-H2O+H]+ | 0.966 | 578.3075562 | 6.306462 | C30H41N7O6 | KUKQXNQRXXTUEM-UHFFFAOYSA-N |
| 4333 | GE20372 B | [M-H2O+H]+ | 0.99 | 594.3027344 | 6.725556 | C30H41N7O7 | DYNPEHYVIZVLIF-LFBFJMOVBZ |
| 4371 | 7-O-methyl-genistein | [M+H]+ | 0.985 | 285.0757446 | 6.746909 | C16H12O5 | KQMVAGISDHMXJJ-UHFFFAOYAR |
| 4416 | colibrimycin B1 | [M+H]+ | 0.998 | 364.0691528 | 6.423467 | C16H14N3O5Cl | MIPKNUAVFFCYQD |

| | | | | | | | |
|---|---|---|---|---|---|---|---|
| 4460 | Chymostatinol_A | [2M+H]+ | 0.998 | 1191.631836 | 6.483975 | C30H41N7O6 | KUKQXNQRXXTUEM-UHFFFAOYSA-N |
| 4463 | Chymostatinol_A | [M+H]+ | 0.99 | 596.3188782 | 6.631195 | C30H41N7O6 | KUKQXNQRXXTUEM-UHFFFAOYSA-N |
| 4580 | Daidzein | [M+H]+ | 0.994 | 255.0651474 | 6.5621395 | C15H10O4 | ZQSIJRDFPHDXIC-UHFFFAOYAG |
| 4583 | Daidzein | [M+Na]+ | 0.982 | 277.0468445 | 6.7177706 | C15H10O4 | ZQSIJRDFPHDXIC-UHFFFAOYAG |
| 4589 | Chymostatinol_A | [M+Na]+ | 0.976 | 618.300415 | 6.586691 | C30H41N7O6 | KUKQXNQRXXTUEM-UHFFFAOYSA-N |
| 4599 | GE20372 B | [M-H2O+H]+ | 0.986 | 594.3027344 | 6.725556 | C30H41N7O7 | DYNPEHYVIZVLIF-LFBFJMOVBZ |
| 4602 | Abyssomicin D | [M+H]+ | 0.866 | 349.1644897 | 6.7938294 | C19H24O6 | BGHABCOWYJWZFY |
| 4678 | CHYMOSTATINOL_C | [M-H2O+H]+ | 0.989 | 592.3237305 | 6.658875 | C31H43N7O6 | WWGFZZYPXWHLAT |
| 4693 | TPU-0037-A_Lydicamycin | [M+Na]+ | 0.976 | 863.5137329 | 6.8213067 | C46H72N4O10 | UXOJELXNKQYYOV-HGMMWVQVSA-N |
| 4710 | colibrimycin A1 demethyl-analogue | [M+H]+ | 0.988 | 574.1691284 | 6.7225056 | C26H28N5O8Cl | NVTOXEPKCFHQMU |
| 4724 | TPU-0037-A_Lydicamycin | [M+H]+ | 0.982 | 841.5321859 | 6.7361765 | C46H72N4O10 | UXOJELXNKQYYOV-HGMMWVQVSA-N |
| 4733 | 7-O-methyl-genistein | [M+H]+ | 0.993 | 285.0757446 | 6.746909 | C16H12O5 | KQMVAGISDHMXJJ-UHFFFAOYAR |
| 4768 | NAI-107_F1 | [M+2H]+2 | 0.998 | 1107.911906 | 6.8060794 | C94H128N26O27S5 | MAVORMFYSNICDR |
| 4832 | NAI-107_F2 | [M+2H]+2 | 0.877 | 1099.412354 | 6.900795 | C94H128N26O26S5 | VDFSODDEUUKYRT |
| 4895 | TPU-0037-B | [M+2H]+2 | 0.86 | 419.271759 | 6.859396 | C47H72N4O9 | PIZUDZCNPHHDQF-CVRRLVMLBC |
| 4904 | Lydicamycin | [M+H]+ | 0.983 | 855.5470581 | 6.9443917 | C47H74N4O10 | UYUJQZZZNYPGMI-JWEMKCLWBK |
| 4923 | Chymostatinol_A | [M+H]+ | 0.984 | 596.3188782 | 6.631195 | C30H41N7O6 | KUKQXNQRXXTUEM-UHFFFAOYSA-N |
| 5133 | Indolokine_A4 | [M+H]+ | 0.959 | 275.0485687 | 7.226674 | C13H10N2O3S | BUZANEDWSLELSW-SNVBAGLBSA-N |
| 5176 | NAI-107_A1 | [M+2H]+2 | 0.999 | 1125.39386 | 7.276786 | C94H127N26O27ClS5 | JZGNTPFPGXVQKP |
| 5217 | NAI-107_A2 | [M+2H]+2 | 0.999 | 1117.394613 | 7.1472087 | C94H127N26O26ClS5 | CEGKQFIOBQHXLN |
| 5272 | CHYMOSTATINOL_C | [M+2H]+2 | 0.905 | 305.6708984 | 7.277895 | C31H43N7O6 | WWGFZZYPXWHLAT |
| 5279 | colibrimycin A1 demethyl-analogue | [M+Na]+ | 0.996 | 596.1513062 | 7.3024907 | C26H28N5O8Cl | NVTOXEPKCFHQMU |
| 5284 | CHYMOSTATINOL_C | [M+H]+ | 0.995 | 610.3343506 | 7.336743 | C31H43N7O6 | WWGFZZYPXWHLAT |

| | | | | | | | |
|---|---|---|---|---|---|---|---|
| 5304 | colibrimycin A1 demethyl-analogue | [M-H2O+H]+ | 1 | 556.1585693 | 7.272173 | C26H28N5O8Cl | NVTOXEPKCFHQMU |
| 5305 | colibrimycin A1 demethyl-analogue | [M+H]+ | 0.987 | 574.1690674 | 7.240354 | C26H28N5O8Cl | NVTOXEPKCFHQMU |
| 5330 | colibrimycin A1 demethyl-analogue | [M+NH4]+ | 1 | 591.1955566 | 7.28731 | C26H28N5O8Cl | NVTOXEPKCFHQMU |
| 5334 | colibrimycin A1 demethyl-analogue | [2M+NH4]+ | 0.992 | 1164.359009 | 7.352495 | C26H28N5O8Cl | NVTOXEPKCFHQMU |
| 5354 | Chymostatin_analog_4 | [M+H]+ | 0.993 | 624.3137207 | 7.040527 | C31H41N7O7 | FXFCCWNGVBFNQW-UHFFFAOYSA-N |
| 5364 | NAI-107_A0 | [M+2H]+2 | 1 | 1108.897705 | 7.339111 | C94H127N26O25ClS5 | RGGRUCPBWQZUOH |
| 5456 | TPU-0037-A_Lydicamycin | [M-H2O+H]+ | 0.984 | 823.5205688 | 7.402214 | C46H72N4O10 | UXOJELXNKQYYOV-HGMMWVQVSA-N |
| 5511 | Genistein | [M+H]+ | 0.993 | 271.06014 | 7.48228 | C15H10O5 | TZBJGXHYKVUXJN-UHFFFAOYAH |
| 5519 | colibrimycin A1 | [2M+NH4]+ | 1 | 1192.391724 | 7.644492 | C27H30N5O8Cl | GIVNVUZVNHSQTA |
| 5538 | colibrimycin A1 | [M+Na]+ | 1 | 610.1664429 | 7.4601126 | C27H30N5O8Cl | GIVNVUZVNHSQTA |
| 5563 | Heptapropylene glycol | [M+NH4]+ | 0.987 | 442.3372192 | 7.4934664 | C21H44O8 | OWRNLGZKEZSHGO |
| 5569 | Furaquinocin_I | [M+H]+ | 0.814 | 417.1542664 | 7.8462915 | C22H24O8 | JJZAZDAEVNRFGT-RMKNXTFCSA-N |
| 5576 | colibrimycin A1 | [M-H2O+H]+ | 1 | 570.1740112 | 7.503847 | C27H30N5O8Cl | GIVNVUZVNHSQTA |
| 5578 | colibrimycin A1 | [M+H]+ | 1 | 588.1845093 | 7.5647154 | C27H30N5O8Cl | GIVNVUZVNHSQTA |
| 5581 | colibrimycin A1 | [2M+H]+ | 1 | 1175.363403 | 7.6125293 | C27H30N5O8Cl | GIVNVUZVNHSQTA |
| 5588 | indolokine_A5 | [M+H]+ | 0.837 | 273.0327148 | 7.640053 | C13H8N2O3S | FWXYUFQKPJMQEC-UHFFFAOYSA-N |
| 5598 | colibrimycin A1 | [M+NH4]+ | 1 | 605.2111206 | 7.5198975 | C27H30N5O8Cl | GIVNVUZVNHSQTA |
| 5610 | TPU-0037-C | [M+H]+ | 0.992 | 825.5365601 | 7.6355953 | C46H72N4O9 | ABOXJBHGJDKUNW-HUVOPICPBK |
| 5612 | TPU-0037-C | [M+Na]+ | 0.979 | 847.5188599 | 7.5067573 | C46H72N4O9 | ABOXJBHGJDKUNW-HUVOPICPBK |
| 5625 | NAI-107_B2 | [M+2H]+2 | 0.997 | 1124.896118 | 7.5392118 | C94H127N26O27ClS5 | WLACVJIBHWIVAC |
| 5637 | TPU-0037-B | [M+H]+ | 0.994 | 837.5369873 | 7.6653852 | C47H72N4O9 | PIZUDZCNPHHDQF-CVRRLVMLBC |
| 5638 | TPU-0037-B | [M+Na]+ | 0.977 | 859.5189209 | 7.595771 | C47H72N4O9 | PIZUDZCNPHHDQF-CVRRLVMLBC |
| 5779 | Propeptin | [M+2H]+2 | 0.937 | 1149.036377 | 7.8542786 | C113H142N26O27 | VIRBPQIAVNKZST |

| | | | | | | | |
|---|---|---|---|---|---|---|---|
| 5907 | Dactylosporolide_A | [M+H]+ | 0.987 | 825.4970703 | 7.7804675 | C42H74O14 | LBOPXMFKHNOMCP |
| 5962 | TPU-0037-B | [M+H]+ | 0.962 | 837.5369873 | 7.6653852 | C47H72N4O9 | PIZUDZCNPHHDQF-CVRRLVMLBC |
| 6020 | 5,6-Dihydroxyphenazin-1(5H)-one | [M+H]+ | 0.988 | 229.0605469 | 8.238041 | C12H8N2O3 | ZXBAHZGKYRGBPG |
| 6040 | Azalomycin F4a | [M+H]+ | 0.988 | 1082.671753 | 8.502236 | C56H95N3O17 | BQEFTEXUBGKSRY-OQEYWIFFSA-N |
| 6046 | Surugamide_F | [M+H]+ | 0.986 | 1056.643921 | 8.217416 | C52H85N11O12 | GDDHFKJRLIQMQR-OJFVTMHKBF |
| 6047 | PHENAZINE | [M+H]+ | 0.987 | 181.0758972 | 8.160814 | C12H8N2 | PCNDJXKNXGMECE |
| 6052 | Surugamide_F | [M+2H]+2 | 0.94 | 528.8261108 | 8.126271 | C52H85N11O12 | GDDHFKJRLIQMQR-OJFVTMHKBF |
| 6089 | "1-Hydroxy-6-methoxy-phenazine;_6-Methoxy-1-phenazinol" | [M+H]+ | 0.979 | 227.0813751 | 8.319198 | C13H10N2O2 | DUXXRWZHWRRFTL |
| 6133 | Levorin A3 | [M+H]+ | 0.894 | 1111.59314 | 8.310894 | C59H86N2O18 | BVEQUWMYNKLPDF-WCHVHIMMSA-N |
| 6177 | SOYASAPONINE_I | [M+H]+ | 0.976 | 943.5249634 | 8.386389 | C48H78O18 | HAQYPCUZKKYSEF |
| 6183 | N-Acetyl-Acyl-Desferrioxamine_C9 | [M+K]+ | 0.973 | 753.4515381 | 8.481658 | C35H66N6O9 | ZVGAZACXCDJNRX |
| 6193 | SOYASAPONINE_I | [M+Na]+ | 0.998 | 965.5078735 | 8.3838 | C48H78O18 | HAQYPCUZKKYSEF |
| 6210 | Levorin A2 | [M+H]+ | 0.97 | 1109.579468 | 8.460233 | C59H84N2O18 | OPGSFDUODIJJGF-WCHVHIMMSA-N |
| 6226 | N-Acetyl-Acyl-Desferrioxamine_C9 | [M+2H]+2 | 0.949 | 358.2515869 | 8.5134325 | C35H66N6O9 | ZVGAZACXCDJNRX |
| 6231 | N-Acetyl-Acyl-Desferrioxamine_C9 | [M+H]+ | 0.981 | 715.4962769 | 8.488495 | C35H66N6O9 | ZVGAZACXCDJNRX |
| 6234 | N-Acetyl-Acyl-Desferrioxamine_C9 | [M+Na]+ | 0.996 | 737.4777222 | 8.393878 | C35H66N6O9 | ZVGAZACXCDJNRX |
| 6316 | Azalomycin F3a | [M+H]+ | 0.993 | 1068.65686 | 8.641783 | C55H93N3O17 | UVUPYXTUQSCQRV-DFIBHIQRSA-N |
| 6371 | COPROPORPHYRIN_III | [M+2H]+2 | 0.98 | 328.1415405 | 8.645193 | C36H38N4O8 | JWFCYWSMNRLXLX |
| 6378 | COPROPORPHYRIN_III | [M+H]+ | 0.988 | 655.2758179 | 8.647747 | C36H38N4O8 | JWFCYWSMNRLXLX |
| 6410 | Azalomycin F4a | [M+H]+ | 0.986 | 1082.671753 | 8.502236 | C56H95N3O17 | BQEFTEXUBGKSRY-OQEYWIFFSA-N |
| 6416 | 1,6-Dihydroxyphenazine | [M+H]+ | 0.99 | 213.0657501 | 8.671864 | C12H8N2O2 | JOXNFMAXWAPITK |
| 6455 | Azalomycin F4a | [M+H+Na]+2 | 0.963 | 552.8309326 | 8.795792 | C56H95N3O17 | BQEFTEXUBGKSRY-OQEYWIFFSA-N |

| | | | | | | | |
|---|---|---|---|---|---|---|---|
| 6482 | Siomycin A | [M+2H]+2 | 0.861 | 824.7371826 | 8.748048 | C71H81N19O18S5 | AKFVOKPQHFBYCA-UHFFFAOYSA-N |
| 6493 | AZALOMYCIN_F-5A | [M+H]+ | 0.963 | 1096.688843 | 9.229502 | C57H97N3O17 | ULHJRSBSTUGEPH |
| 6525 | colibrimycin C6 | [M+H]+ | 1 | 430.1523743 | 8.890668 | C22H24N3O4Cl | FOJYTDOWSIXHIB |
| 6527 | colibrimycin C6 | [M+Na]+ | 0.995 | 452.1348267 | 8.718899 | C22H24N3O4Cl | FOJYTDOWSIXHIB |
| 6552 | Surugamide_A | [2M+H]+ | 0.987 | 1824.251587 | 8.744501 | C48H81N9O8 | NPYICXUUGUJPMM-UHFFFAOYSA-N |
| 6557 | Champacyclin | [M+2H]+2 | 0.952 | 456.8173828 | 8.931428 | C48H81N9O8 | ARNLSFQSZWTPRF-YOHDJLIIBL |
| 6558 | Surugamide_A | [M+H]+ | 0.991 | 912.62677 | 8.972838 | C48H81N9O8 | NPYICXUUGUJPMM-UHFFFAOYSA-N |
| 6559 | Surugamide_A | [M+K]+ | 0.981 | 950.5824585 | 8.815069 | C48H81N9O8 | NPYICXUUGUJPMM-UHFFFAOYSA-N |
| 6566 | Champacyclin | [M+Na]+ | 0.988 | 934.6092529 | 8.786918 | C48H81N9O8 | ARNLSFQSZWTPRF-YOHDJLIIBL |
| 6631 | Acyl-Desferrioxamine_C12 | [M+Na]+ | 0.995 | 737.5134277 | 8.893016 | C36H70N6O8 | MMADSZAPNXUIPH |
| 6651 | Acyl-Desferrioxamine_C12 | [M+H]+ | 0.999 | 715.5327148 | 9.111518 | C36H70N6O8 | MMADSZAPNXUIPH |
| 6652 | Acyl-Desferrioxamine_C12 | [M+2H]+2 | 0.973 | 358.2695007 | 8.890727 | C36H70N6O8 | MMADSZAPNXUIPH |
| 6767 | Azalomycin F5a | [M+H]+ | 0.986 | 1096.688965 | 9.363718 | C57H97N3O17 | SPNDCJQGKUZBHK-HQKIQJKYSA-N |
| 6895 | Alteramide A | [2M+Na]+ | 0.881 | 1043.533936 | 9.8764715 | C29H38N2O6 | HZKFYHKCBBWPIX-HKZOSMIOSA-N |
| 6901 | Alteramide A | [M+H]+ | 0.973 | 511.2793274 | 9.75102 | C29H38N2O6 | HZKFYHKCBBWPIX-HKZOSMIOSA-N |
| 6906 | Alteramide A | [2M+H]+ | 0.891 | 1021.551819 | 9.568613 | C29H38N2O6 | HZKFYHKCBBWPIX-HKZOSMIOSA-N |
| 6918 | Alteramide A | [M+Na]+ | 0.985 | 533.2616577 | 9.7719755 | C29H38N2O6 | HZKFYHKCBBWPIX-HKZOSMIOSA-N |
| 7158 | Alteramide B | [M+H]+ | 0.983 | 495.2846222 | 10.363046 | C29H38N2O5 | RKVKDOPZGFNWBV-CTTZNNTJSA-N |
| 7160 | Ikarugamycin_epoxide | [2M+H]+ | 0.959 | 989.5622559 | 10.226913 | C29H38N2O5 | SGDSWAQEEGMGDC-UHFFFAOYSA-N |
| 7334 | Maltophilin | [M-H2O+H]+ | 0.969 | 493.2690735 | 11.173563 | C29H38N2O6 | ZIFOVGPOLIICJX-FDCDIUKQSA-N |
| 7365 | MARINOMYCIN_A | [M-H2O+H]+ | 1 | 979.5200195 | 10.995573 | C58H76O14 | GXBIDOSKEKEFEF-UHFFFAOYSA-N |

| | | | | | | | |
|---|---|---|---|---|---|---|---|
| 7366 | MARINOMYCIN_A | [M+H]+ | 0.807 | 997.529541 | 11.001315 | C58H76O14 | GXBIDOSKEKEFEF-UHFFFAOYSA-N |
| 7399 | Antimycin A5 | [M+H]+ | 0.921 | 493.2178955 | 11.276923 | C24H32N2O9 | QVLIJBABYDIICV-UHFFFAOYSA-N |
| 7411 | MARINOMYCIN_A | [M+NH4]+ | 0.946 | 1014.557007 | 11.508205 | C58H76O14 | GXBIDOSKEKEFEF-UHFFFAOYSA-N |
| 7528 | Antimycin A20 | [M+H]+ | 0.825 | 507.2330017 | 11.656901 | C25H34N2O9 | JFLKVAOGXZTMRU-LKMKPSOZSA-N |
| 7738 | Antimycin A17 | [M+H]+ | 0.937 | 535.2643229 | 12.477216 | C27H38N2O9 | CRIAYTZRPXXKCY-UHFFFAOYSA-N |
| 7775 | PIERICIDIN_A1 | [M+H]+ | 0.982 | 416.2793884 | 12.789845 | C25H37NO4 | BBLGCDSLCDDALX |
| 7783 | PIERICIDIN_A1 | [M-H2O+H]+ | 0.982 | 398.2686462 | 12.716675 | C25H37NO4 | BBLGCDSLCDDALX |
| 7840 | Antimycin A19 | [M+Na]+ | 0.987 | 571.2627563 | 12.967299 | C28H40N2O9 | FKJLKGUXRIDXTL-DQRKQOBLSA-N |
| 7903 | 21-hydroxy oligomycin C | [M-H2O+H]+ | 0.957 | 773.5198975 | 13.054773 | C45H74O11 | PDHNJYCZSJPJDI-XHTHQMERBX |
| 7904 | 21-hydroxy oligomycin C | [M+H]+ | 0.912 | 791.530599 | 13.092589 | C45H74O11 | PDHNJYCZSJPJDI-XHTHQMERBX |
| 7908 | Nigericin | [M-H2O+H]+ | 0.995 | 707.4725952 | 13.032956 | C40H68O11 | DANUORFCFTYTSZ-GVMXEALNSA-N |
| 7909 | Nigericin | [M+NH4]+ | 0.988 | 742.5102539 | 13.143413 | C40H68O11 | DANUORFCFTYTSZ-GVMXEALNSA-N |
| 7945 | Antimycin A15 | [M+H]+ | 0.869 | 563.2958374 | 13.370498 | C29H42N2O9 | JRUNAPYTQMZOTG-UHFFFAOYSA-N |
| 8337 | Ornithine-containing lipid_597 | [M+H]+ | 0.991 | 597.5199992 | 14.884098 | C35H68N2O5 | ANJLTOOLWORDTB-UHFFFAOYSA-N |
| 8359 | S-3466 A | [M+NH4]+ | 0.966 | 782.5057983 | 14.802821 | C42H68O12 | ZBDGIMZKOJALMU-UHFFFAOYSA-N |
| 8412 | Ornithine-containing lipid-like_611 | [M+H]+ | 0.983 | 611.5357869 | 14.973755 | C36H70N2O5 | ANJLTOOLWORDTB-UHFFFAOYSA-N |
| 8444 | S-3466 B | [M+NH4]+ | 0.985 | 796.5211182 | 14.830408 | C43H70O12 | DFQMKYUSAALDDY-UHFFFAOYSA-N |
| 8452 | Ornithine-containing-lipid-like | [M+ACN+H]+ | 0.991 | 639.5668335 | 15.065583 | C38H74N2O5 | ANJLTOOLWORDTB-UHFFFAOYSA-N |
| 8481 | Nigericin | [M+K]+ | 0.982 | 763.4395142 | 14.783022 | C40H68O11 | DANUORFCFTYTSZ |
| 8491 | TETRANACTIN | [M+Na]+ | 0.989 | 815.4907837 | 14.859801 | C44H72O12 | NKNPHSJWQZXWIX-UHFFFAOYSA-N |
| 8500 | TETRANACTIN | [M+NH4]+ | 0.994 | 810.5358887 | 15.219181 | C44H72O12 | NKNPHSJWQZXWIX-UHFFFAOYSA-N |
| 8514 | Nigericin | [M-H2O+H]+ | 1 | 707.4729004 | 14.846936 | C40H68O11 | DANUORFCFTYTSZ |

| | | | | | | | |
|---|---|---|---|---|---|---|---|
| 8516 | Nigericin | [M+NH4]+ | 0.989 | 742.510376 | 14.871661 | C40H68O11 | DANUORFCFTYTSZ |
| 8535 | Ornithine-containing lipid_597 | [M+H]+ | 0.988 | 597.5199992 | 14.884098 | C35H68N2O5 | ANJLTOOLWORDTB-UHFFFAOYSA-N |
| 8538 | Ornithine-containing lipid | [M+H]+ | 0.99 | 625.551534 | 15.260318 | C37H72N2O5 | PUZMSNPCMYUOTE-UHFFFAOYSA-N |
| 8546 | Ornithine-containing-lipid-like | [M+Na]+ | 0.99 | 661.5487671 | 14.59718 | C38H74N2O5 | ANJLTOOLWORDTB-UHFFFAOYSA-N |
| 8547 | Ornithine-containing-lipid-like | [2M+H]+ | 0.898 | 1278.125854 | 15.236974 | C38H74N2O5 | ANJLTOOLWORDTB-UHFFFAOYSA-N |
| 8555 | Ornithine-containing lipid-like_611 | [M+H]+ | 0.99 | 611.5357869 | 14.973755 | C36H70N2O5 | ANJLTOOLWORDTB-UHFFFAOYSA-N |
| 8584 | Pamamycin-691 | [M+K]+ | 0.92 | 730.5020142 | 14.973916 | C41H73NO7 | CHIITYMJWGPWLG |
| 8637 | Ornithine-containing-lipid-like | [M+ACN+H]+ | 0.993 | 639.5668335 | 15.065583 | C38H74N2O5 | ANJLTOOLWORDTB-UHFFFAOYSA-N |
| 8702 | Ornithine-containing lipid_653 | [M+Na]+ | 0.975 | 675.5637817 | 14.828809 | C39H76N2O5 | DWOURMRLOILMHL-UHFFFAOYSA-N |
| 8703 | Ornithine-containing lipid_653 | [2M+H]+ | 0.961 | 1306.157349 | 14.840947 | C39H76N2O5 | DWOURMRLOILMHL-UHFFFAOYSA-N |
| 8721 | Nigericin like | [M-H2O+H]+ | 0.919 | 691.4782715 | 15.062841 | C40H68O10 | ANJLTOOLWORDTB-UHFFFAOYSA-N |
| 8762 | Ornithine-containing lipid | [M+H]+ | 0.989 | 625.551534 | 15.260318 | C37H72N2O5 | PUZMSNPCMYUOTE-UHFFFAOYSA-N |
| 8829 | WA8242A1 | [M+H]+ | 0.907 | 683.5571289 | 15.424381 | C39H74N2O7 | BSNMYXPQSLGSDY-UHFFFAOYSA-N |
| 8916 | Ornithine-containing-lipid-like | [M+ACN+H]+ | 0.992 | 639.5668335 | 15.065583 | C38H74N2O5 | ANJLTOOLWORDTB-UHFFFAOYSA-N |
| 9061 | Ornithine-containing lipid_653 | [M+H]+ | 0.96 | 653.5827433 | 15.659049 | C39H76N2O5 | DWOURMRLOILMHL-UHFFFAOYSA-N |
| 9211 | WA8242A1 | [M+H]+ | 0.927 | 683.5571289 | 15.424381 | C39H74N2O7 | BSNMYXPQSLGSDY-UHFFFAOYSA-N |
| 9367 | Ornithine-containing lipid | [M-H2O+H]+ | 0.97 | 607.5403442 | 15.824362 | C37H72N2O5 | PUZMSNPCMYUOTE-UHFFFAOYSA-N |
